# Brain *Mecp2* Gene Dosage and Gene Therapy Shape Multi-Omic Signatures and Putative Biomarkers in Rett Syndrome

**DOI:** 10.1101/2025.08.30.673242

**Authors:** Stephanie A. Zlatic, Eric Dammer, Amanda Crocker, Duc Duong, Jim Selfridge, Kamal KE Gadalla, Avanti Gokhale, Brendan R. Tobin, Levi B. Wood, Martina Zandl-Lang, Lucia Abela, Barbara Plecko, Anupam Patgiri, Walter E. Kaufmann, Randall Carpenter, Stuart Cobb, Victor Faundez

**Author notes:** These authors contributed equally (Stephanie A. Zlatic, Eric Dammer, Amanda Crocker, Duc Duong). Department of Biosciences, Nottingham Trent University; Nottingham, NG11 8NS, UK.

## Abstract

Rett syndrome (RTT) is a neurodevelopmental disorder caused by *MECP2* mutations. Like other genetic neurodevelopmental disorders, it lacks protein biomarkers to evaluate disease and therapeutic outcomes. We present a strategy to define putative biomarkers of MeCP2 dysfunction in brain with potential to delineate mechanisms and monitor therapeutic interventions. This strategy relies on a library of proteins responsive to *Mecp2* gene dosage and correlated with molecular and clinical outcomes after AAV9-mediated *MECP2* gene therapy in *Mecp2*-KO mice. Gene rescue restored MeCP2 in brain, improved clinical phenotypes, and reverted transcriptome and proteome abnormalities. We identified 327 shared proteins among 1852 cortical and hippocampal proteins responsive to *Mecp2*/*MECP2*. Of these, 119 also displayed Mecp2/MECP2-dependent transcript changes. Both the Mecp2-responsive proteome and transcript–protein pairs were enriched in synaptic and metabolic pathways, including central carbon and NAD+ metabolism. We used this therapy-responsive protein library to guide selection of candidate cerebrospinal fluid (CSF) biomarkers in RTT. CSF composition from neurotypical and RTT groups was analyzed using ultrasensitive nucleic acid-based multiplexed ELISA. Twenty-eight proteins were altered in RTT, nine overlapping with *Mecp2* dosage- and therapy-sensitive proteins. Multivariate regression linked several candidates to *Mecp2*/*MeCP2* abundance and phenotypic improvement in mice. This paradigm provides a rigorous molecular systems-level framework integrating genetics, preclinical gene therapy, and clinical metrics to define robust cross-species putative biomarkers and possible mechanisms in RTT, with potential applicability to other neurodevelopmental disorders.

**One Sentence Summary:** Genetic Identification of cross-species biomarkers and mechanisms in Rett Syndrome

## Introduction

The cellular and molecular understanding of neurodevelopmental disorders is advancing through the study of monogenic syndromes modeled in preclinical mouse models. Among these, Rett syndrome (RTT) stands out due to its profound and diverse neurological impairments, developmental regression, and the availability of multiple well-characterized murine disease models (*1–3*). Rett syndrome is preferentially caused by mutations in the *MECP2* gene, which encodes MeCP2, an epigenetic factor and RNA polymerase interactor that modulates the transcription of numerous RNAs in the brain (*1, 4, 5*). MeCP2 binds to nearly 80% of neuronal chromatin, orchestrating complex transcriptomic programs across diverse brain regions and cell types (*6–11*). While RTT is monogenic, the breadth of genes affected by MeCP2 dysfunction renders the syndrome functionally akin to a polygenic disorder. This molecular complexity complicates the identification of biomarkers of disease and pathogenic mechanisms, which currently include synaptic and metabolic defects (*1, 2, 12*). The relative contribution and hierarchy of these mechanisms in shaping disease are under active investigation.

Defining prioritized disease mechanisms and their molecular components is essential for the development of biomarkers. Extensive transcriptomic studies in preclinical models and samples from individuals with RTT have identified reproducible MeCP2-responsive transcripts with strong potential as molecular biomarkers (*1, 3, 13–15*). However, comparable systematic efforts to identify protein biomarkers, which may more directly reflect disease biology and provide clinically accessible readouts, remain limited (*16–18*). The need for mechanism-based biomarkers, particularly those related to MeCP2 expression, is underscored by recent advances in gene and growth factor therapies, which offer promising avenues for RTT treatment (*1*). The efficacy of these interventions is being assessed using standardized clinical evaluations in both humans and mice (*13, 19, 20*). In the case of RTT, gene therapy success hinges on not only the delivery of *MECP2* but also on achieving precise expression levels, as both deficiency and overexpression of *MECP2* are pathogenic (*1*). As such, molecular biomarkers of MeCP2 dysfunction should dynamically respond across a spectrum ranging from a null allele of *MECP2* to a duplication of the gene.

To address the unmet need for mechanistically informed biomarker discovery, we developed a genetic strategy in preclinical models aimed at enabling cross-species identification of putative biomarkers of brain MeCP2 dysfunction detectable in human biofluids. We defined a library of mouse brain proteins and transcripts based on three stringent criteria. First, candidate molecules had to exhibit coherent and reciprocal changes in response to *Mecp2* loss- and gain-of-function in mice to mimic RTT and *MECP2* duplication syndromes, using *Mecp2* knockout and Tau-driven *Mecp2* overexpression models, respectively (*21, 22*). Second, these dysregulated proteins and transcripts in *Mecp2*-null brains had to be corrected by AAV9-mediated *MECP2* gene therapy (*19*). Third, their expression levels had to correlate with phenotypic improvements in gene therapy–treated mice. We reasoned that molecules meeting these criteria would comprise a genetically and therapeutically validated panel of proteins and inform putative mechanisms to serve as a source for candidate biomarkers in biosamples from individuals with RTT.

To implement this strategy, we performed intracerebroventricular delivery of an AAV9 vector encoding human *MECP2* engineered with the Expression Attenuation via Construct Tuning (EXACT) system, allowing precise restoration of MeCP2 whilst avoiding pathogenic levels of overexpression (*19*). Our goal was to precisely correlate phenotypic clinical scores and MeCP2 protein levels with their transcriptome and proteome in each animal. Using this strategy, we identified 327 proteins shared by the cortex and hippocampus of *Mecp2*-KO mice, among a total of 1,852 proteins responsive to *Mecp2* gene dosage and *MECP2* gene therapy. To evaluate the translational potential of this protein library, we analyzed cerebrospinal fluid from two independent cohorts of matched neurotypical controls and individuals with RTT using a high-sensitivity nucleic acid-based multiplex ELISA platform. We identified 9 CSF proteins that overlapped with the *Mepc2*-sensitive murine proteome and were significantly altered in patients with RTT. Importantly, the abundance of these proteins correlated with molecular and behavioral phenotypes in mice.

In summary, our preclinical genetic strategy establishes a rigorous, systems-level framework that integrates genetic manipulation and phenotypic correlation to define putative interspecies biomarkers and pathogenic mechanisms in RTT. This approach may be extended to other genetic neurodevelopmental disorders, even those with complex or poorly defined molecular architectures.

## Results

Gene therapy rescued MeCP2 levels and partially rescued clinical phenotypes of *Mecp2*-KO (*Mecp2^-/y^*) mice (Fig. 1). Human *MECP2* gene therapy was performed with the EXACT system (AAV9-RTT271) and outcomes were assessed in a large cohort of eight-week-old wild-type, and *Mecp2-KO* mice treated by intracerebroventricular injection at postnatal day 1 with either AAV9 *MECP2* gene therapy (RTT271) or saline control (*19*). We scored clinical outcomes from the same animals we processed for multiomics analysis. Expression of mouse- and human-specific Mecp2/MeCP2 protein in the cortex was confirmed (Fig. 1A, B). *Mecp2-KO* mice receiving either a 1e11 or 3e11 viral genome (vg) dose of AAV9-RTT271 increased human MeCP2 abundance to the same level, corresponding to 60% of the endogenous wild-type MeCP2 protein levels (Fig. 1A). MeCP2 protein expression improved *Mecp2-KO* phenotypes including apnea, open field test, and a clinical aggregated score where four of the six constituent parameters were restored similarly by both doses of AAV9-RTT271 (Fig. 1D-F) (*19, 23*). We therefore pooled both AAV9-RTT271 doses for all subsequent analyses. Clinical aggregated score improvement correlated with MeCP2 protein levels in either cortex or hippocampus as compared to saline-injected wild-type and *Mecp2*-null mice (Fig. 1G). We did not find signs of toxic responses to AAV vectors either at the level of cortical cytokine mRNAs or the plasma levels of these cytokines, thus excluding overt vector-indued inflammatory responses (Fig. 1S1). These results validated our gene therapy cohorts for further genome-wide expression studies.

**Figure 1.**
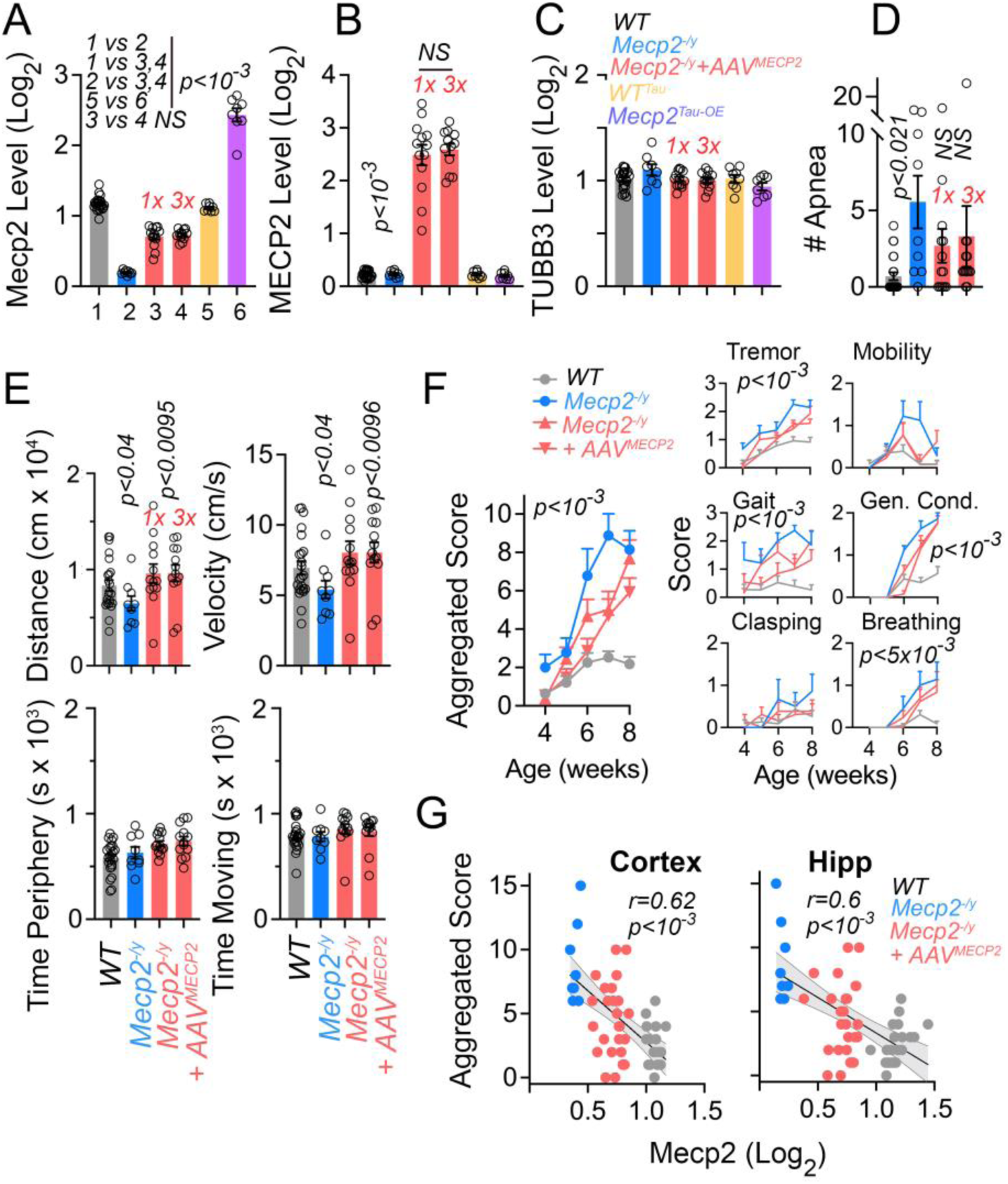
Mouse and Human Mecp2/MeCP2 Protein Expression Levels and Gene Therapy Phenotypic Reversal of *Mecp2*-KO Phenotypes. Mass spectrometry quantification of mouse and human Mecp2/MeCP2 protein with peptides either common to both species (A) or human specific MECP2 peptides (B) and beta tubulin (C, TUBB3), as control in the cortex of wild-type animals (n=23), *Mecp2^-/y^* (n=8), *Mecp2^-/y^* receiving intracerebroventricularly 1e11 AAV9-RTT271 vg per mouse (n=13), *Mecp2^-/y^* receiving 3e11 AAV9-RTT271 vg per mouse (n=12), *Mecp2* overexpression driven from the Tau promoter (n=8) and their wild-type controls (yellow column 5, n=8). 1x and 3x denote the amount of vg delivered intracerebroventricularly at postnatal day 0. D. Number of apneas per 20 seconds in animals of the specified genotype without or with AAV9-RTT271 treatment. A-D, One Way ANOVA followed by Šídák’s multiple comparisons. E. Open field test phenotypes in animals of the specified genotype without or with AAV9-RTT271 treatment. One Way ANOVA followed by Benjamini, Krieger and Yekutieli corrections. F. Aggregated phenotypic score and its constituent parameters in animals of the specified genotype without or with AAV9-RTT271 treatment. Mixed-effect analysis followed by Tukey’s multiple comparisons test, p value depicts the effect of genotype and gene therapy effects on the parameter (Gene and Gene Therapy Effect F (3, 54) = 16.79, Time Effect F (3.482, 181.1) = 51.58, Time x Gene and Gene Therapy Interaction F (12, 208) = 5.108). Inverted triangles represent 3e11 vg dose. G. MeCP2 protein levels in A inversely correlate with the aggregated phenotypic score in F in two brain regions. Least squares non-linear regression and 95% confidence bands in gray.

### Brain Transcriptomic and Proteomic Consequences of *Mecp2* Loss- and Gain-of-Function Alleles and Their Rescue by Gene Therapy

We quantified the coding and small non-coding transcriptomes in the cortex and hippocampus of the experimental groups described above and from mice overexpressing *Mecp2* from the Tau promoter and their isogenic controls. *Mecp2 ^Tau-OE^* mice produce two-fold MeCP2 proteins compared to wild-type levels (Fig. 1A, compare columns 5-6). Both doses of AAV9-RTT271 expressed the same *MECP2* mRNA level in cortex (Fig. 2A, 1x=928±104 counts and 3x=1248±175, p=0.159) while the expression of *Mecp2* driven by the Tau promoter was ∼4-fold of the wild-type *Mecp2* mRNA levels in cortex and hippocampus (Fig. 2A, compare yellow and purple bars, p<0.001). We selected *Mecp2* dosage-sensitive transcriptomes, here defined as “q-defined transcriptomes”, by thresholding datasets with a FDR-corrected multiple ANOVA across all experimental conditions. Our goal was to segregate experimental groups by genotype and AAV9-RTT271 therapy, as determined by principal component analysis and Euclidean distance clustering (Fig. 2B). Coding transcriptomes were the most affected transcriptomes in cortex and hippocampus as determined by percent of all expressed genes (Fig. 2C-E). Cortical coding transcriptome segregated experimental groups at q<0.1 (4736 mRNAs, Fig. 2C). We validated the cortical q-defined coding transcriptome (Fig. 2C) by its similarity with previous cortical *Mecp2*-null coding transcriptomes (r=0.634, Fig. 2S1A, B and D) (*6, 9, 11, 24, 25*). The cortical coding transcriptome was modified by genotype and gene therapy in a coherent manner (Fig. 2C), where coherence is defined as mRNA levels affected in *Mecp2-KO* shifting closer to wild-type levels after gene therapy and changing oppositely in *Mecp2 ^Tau-OE^* as compared to *Mecp2-KO*. The number and magnitude of change in differentially expressed cortical mRNAs in *Mecp2-KO* mice were reduced after AAV9-RTT271 therapy (Fig. 2F). Cortical gene therapy brought the coding transcriptome closer to a wild-type transcriptome, whereas Tau-driven *Mecp2* overexpression significantly modified a discrete set of transcripts (Fig. 2F). Only the q-defined cortical mRNA transcriptome displayed *Mecp2* dose- and gene therapy-coherence across *Mecp2* genotypes and gene therapy conditions as compared to mRNAs in hippocampus as well as all non-coding transcriptomes (Fig. 2C and D). These results define a robust mRNA cortical transcriptome sensitive to gene dosage and reversible by *MECP2* gene therapy.

**Figure 2.**
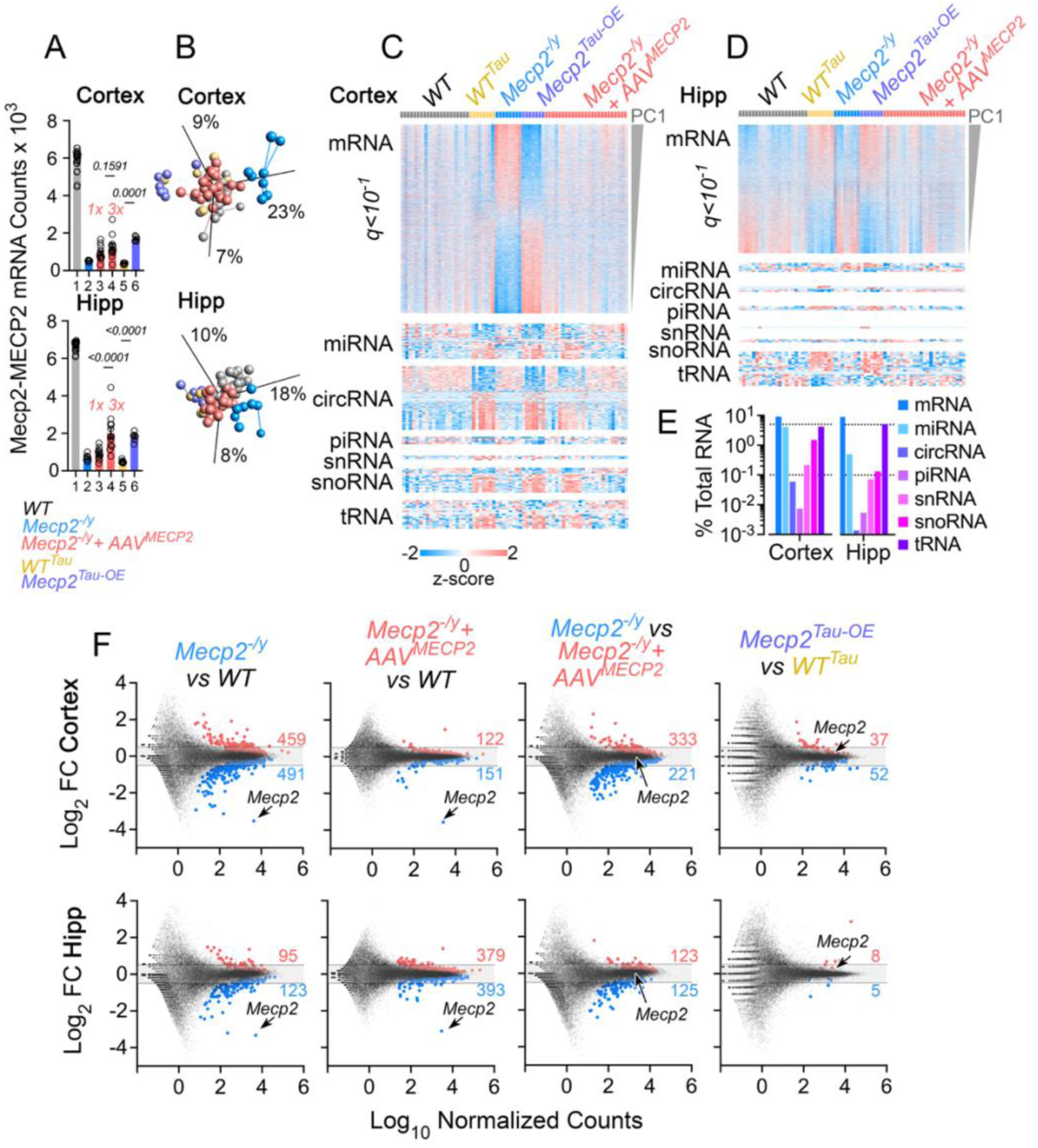
Brain Transcriptomes of *Mecp2^-/y^* and *Mecp2^Tau-OE^* Mice and the Effect of Gene Therapy. A. Normalized counts of mouse *Mecp2* mRNA and human *MECP2* mRNA in the cortex and hippocampus of the animals of the specified genotype without or with AAV9-RTT271 treatment. 1x and 3x denote the amount of vg delivered intracerebroventricularly. Expression was scored with sequences specific to either the endogenous *Mecp2* gene (1–2), the *MECP2* AAV9 sequence (3–4), or Tau-*Mecp2* transgene (5–6). B. Principal component analysis of differentially expressed coding transcriptomes subjected to multiple ANOVA followed by Benjamini-Hochberg FDR correction set at q<0.1 in cortex and hippocampus. Lines among symbols indicate clustered groups defined by Euclidean distance. Colors indicate specified genotype without or with AAV9-RTT271 treatment as in A. C. Cortical Z-scored heat maps of coding and non-coding RNA transcriptomes. Each transcriptome was selected by FDR-corrected multiple ANOVA q<0.1 across all experimental groups. Two doses of AAV9-RTT271 treatment were merged into one category for these analyses. Rows are organized by analyte contribution to PC1. D. Hippocampal Z-scored heat maps of coding and non-coding RNA transcriptomes as in C. E. Percent of all the transcripts quantified in each transcriptome selected after filtering with q<0.1. F. MA plots of coding transcriptomes in cortex and hippocampus comparing genotypes and effects of gene therapy. Red and blue symbols are mRNAs significantly increased or decreased, respectively, in each comparison by Wald test statistics and FDR corrected by Benjamini-Hochberg. Processed data used to build figures can be found in supplementary table I.

Concurrently with the transcriptomes, we measured the cortical and hippocampal proteomes of the wild-type, *Mecp2*-null, *MECP2* gene therapy, and *Mecp2* overexpression mouse coronal sections using Tandem Mass Tagging (*26*). We quantified 10,433 proteins present in cortex and hippocampus which were thresholded by a q value below 0.001 using FDR-corrected multiple ANOVA to select “q-defined proteomes” (Fig. 3A-C). With this threshold, *Mecp2-KO* and *Mecp2* overexpression genotypes segregated away from each other and from a cluster constituted by wild-type and AAV9-RTT271 therapy-treated *Mecp2-KO* animals (Fig. 3A). The q-defined cortical proteome was composed of 1,127 proteins while the q-defined hippocampal proteome comprised 725 proteins (Fig. 3B-C). Proteomes in both brain regions responded coherently to *Mecp2* gene dosage (Fig. 3B-C). Of these cortex and hippocampus *Mecp2*-responsive proteins only 327 were shared between both brain regions (Fig. 3C, 4.2-fold enrichment, p<1E-10 hypergeometric test). Following AAV9-RTT271 treatment, proteome-wide deviations from wild-type were reduced, with the distribution of protein abundance changes converging near the origin, consistent with normalization of the proteome phenotypes in both brain regions (Fig. 3D). Similar proteome-wide attenuation was observed when comparing either wild-type or AAV9-RTT271 therapy-treated animals with *Mecp2-KO* animals (Fig. 3D) while proteome-wide differences were reduced when comparing AAV9-RTT271 therapy-treated with wild-type brain regions (Fig. 3D). These results define cortical and hippocampal proteomes whose robustness relies upon their response to *Mecp2* gene dosage and their restoration by *MECP2* gene therapy. In addition to the coherent proteomes, we identified 900 proteins whose levels were altered in *Mecp2* overexpression mouse cortex (Fig. 3S1). Of these proteins, 200 were only affected in *Mecp2 ^Tau-OE^* cortex thus suggesting unique mechanisms operating in *Mecp2* gain-of-function alleles (Fig. 3S1).

**Figure 3.**
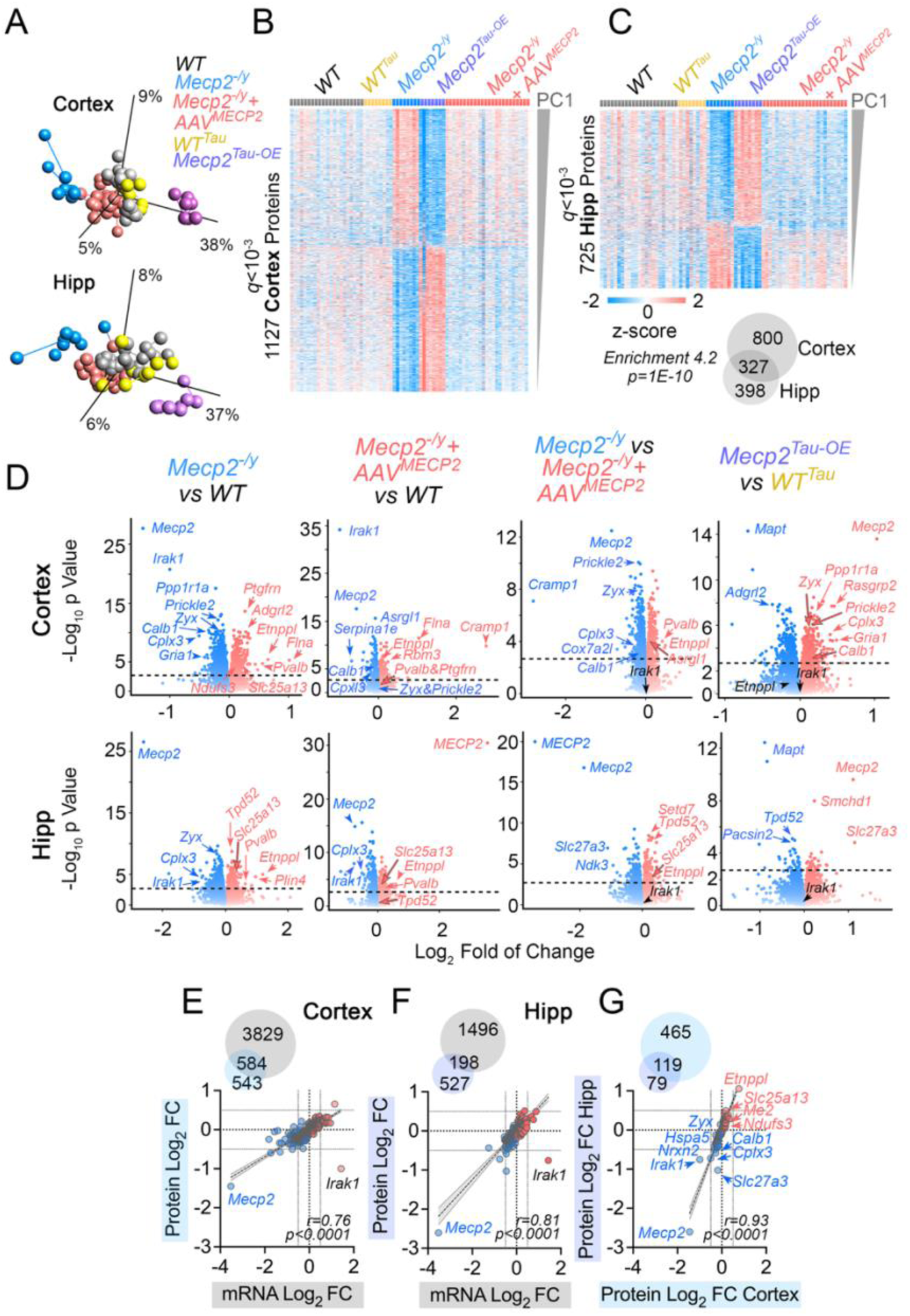
Coherent Brain Proteomes of *Mecp2^-/y^* and *Mecp2^Tau-OE^* Mice and their Restoration by Gene Therapy. A. Principal component analysis of differentially expressed proteins selected by multiple ANOVA followed by Benjamini-Hochberg FDR correction set at q<0.001 in cortex and hippocampus. Lines among symbols indicate clustered groups defined by Euclidean distance. B and C depict cortical and hippocampal Z-scored heat maps of proteomes selected by FDR-corrected multiple ANOVA q<0.001 across all experimental groups. Two doses of AAV9-RTT271 treatment were merged into one category for these analyses. Rows are organized by analyte contribution to PC1. D. Cortical and hippocampal volcano plots comparing genotypes and effects of gene therapy. Red and blue symbols are proteins increased or decreased significantly, respectively, by FDR-corrected t-test set at q<0.001. E. Venn diagram of the 584 overlapping q-defined cortical proteins and mRNAs and their correlation. F. Venn diagram of the 198 overlapping q-defined hippocampal proteins and mRNAs and their correlation. G. Venn diagram of the 584 and 198 q-defined cortical and hippocampal protein-mRNA pairs, in E and F respectively. These protein-mRNA pairs are strongly correlated between the two brain regions. E to G Correlation and p value were obtained by least squares non-linear regression. Depicted are 95% confidence bands in gray.

A total of 584 gene products overlapped between the cortical q-defined proteins and q-defined coding transcriptome (Fig. 3E, mRNA-protein overlap enrichment=2.4, p=5.9e-119, hypergeometric test). This overlap encompasses 51.8% of the cortical proteome yet only 12.2% of the cortical mRNAs sensitive to *Mecp2* gene dosage. Cortical protein and mRNA differential expression changes due to *Mecp2* gene dosage modifications were strongly correlated (r=0.76, Fig. 3E), suggesting these gene products are likely under control of Mecp2 transcriptional regulatory activity. Similar analysis of the hippocampus revealed 198 q-defined proteins whose differential expression levels strongly correlated with their mRNA levels (mRNA-protein overlap enrichment=2.7, p=1.4e-40, hypergeometric test. r=0.81, Fig. 3F). The cortex and hippocampus *Mecp2*-sensitive mRNA-protein pairs converged on 119 strongly correlated protein-transcript pairs (protein overlap enrichment=21.2, p=1.56e-134, hypergeometric test. r=0.91, Fig. 3G). These 119 protein-transcript pairs are enriched in proteins annotated to ontologies of synaptic, mitochondrial, central carbon, and cholesterol metabolism (*24, 27*). (Fig. 3S2)(*28*) and long genes, the latter a feature of direct *Mecp2* target genes (Fig. 2S1G) (*6, 7*). These findings uncover a group of genes whose mRNAs and proteins respond in a similar fashion to *Mecp2* dosage and/or *MECP2* gene therapy irrespective of their anatomical location.

We assessed the reproducibility of these selected 119 protein-mRNA pairs by comparing their expression with published cortical transcriptomes obtained from animals carrying the same *Mecp2* mutation or a mouse model of the MECP2 duplication syndrome (Fig. 4 and 2S1 B-D) (*6, 9, 25*). The expression of these 119 genes robustly and positively correlated in *Mecp2* KO transcriptomes with correlation coefficients ranging from 0.53 to 0.74 (Fig. 2S1B and D) (*6, 9, 25*). In contrast, the correlation was inverted in the transcriptome of a MECP2 duplication syndrome mouse (r=-0.74, Fig. 2S1C) (*29*). This inverse concordance indicates that genes increased in the duplication model are generally decreased in our loss-of-function datasets, and vice versa, consistent with direct regulation by *Mecp2* dosage. Furthermore, the overlap of the 119 protein-mRNA pairs extended to a similar *Mecp2* deficient cortical proteome (*16*). We identified 11 proteins shared between these two datasets whose levels strongly correlated (protein overlap enrichment=8, p=1.43e-7, hypergeometric test. r=0.94, Fig. 2S2).

**Figure 4.**
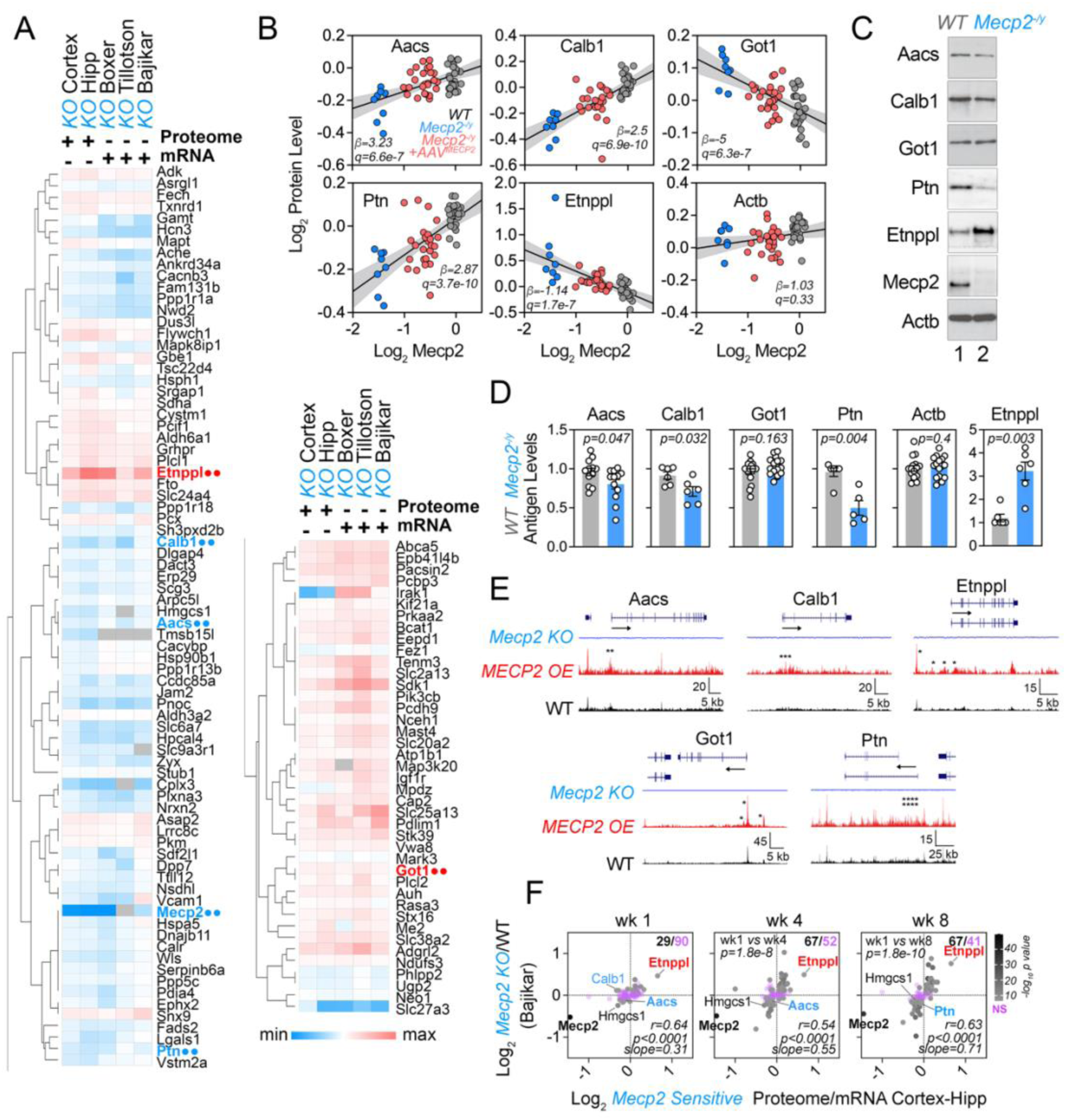
Validation of a Robust Set of 119 *Mecp2-*sensitive Protein-mRNA Pairs Shared Across Brain Regions. A. Z-scored heat map comparing the 119 protein-mRNA pairs selected from cortex and hippocampus depicted and the transcriptome datasets by Boxer et al, Tillotson et al, and Bajikar et al. B. Proteins selected from the panel A by their fold of change across datasets depicted as a correlation with MeCP2 protein levels in cortex. Beta (effect size) corresponding to the slope of a fit line for each protein as a predictor of MeCP2 protein content. Color annotates genotype and gene therapy. Depicted is best linear fit ±95% CI. C. Immunoblot analysis of proteins selected in B. D. Quantification of antigens presented in C. Each dot is an independent cortex. Unpaired median difference two-sided permutation t-test. E. Analysis of Bajikar et al. (2023) CUT&RUN profiling of MeCP2 binding. Wild-type (gray), *Mecp2^-/+^* (blue), and MECP2 OE transgenic mice (red) tracks at the indicated loci. Asterisks mark peaks with significant differences across genotypes located near the gene translation initiation site (see table I for values) Tracks are displayed as an aggregate of biological replicates (n=3–6 animals per genotype). F. Temporal evolution of the mRNA levels after acute deletion of the Mecp2 gene and their correlation with 119 *Mecp2* mutant- and gene therapy-sensitive mRNA-Protein Pairs (data from Bajikar et al 2015). Upper right corner numbers denote in black significantly changed mRNA in the Bajikar dataset and non-significant mRNA (purple). P values between weeks compare number of significantly changed mRNA (Exact binomial McNemar p-value). Pearson correlation coefficient.

The translatability of putative biomarkers prioritized from preclinical animal models depends on how fast these proteins respond to *Mecp2* gene dosage manipulation. We explored how the 119 Mecp2-sensitive proteins respond to acute deletion of the Mecp2 gene using transcriptome datasets after acute *Mecp2* deletion by Bajikar et al (*25*). We first focused on Accs, Calb1, Got1, Ptn, and Etnppl, for orthogonal validation. These proteins undergo some of the most pronounced changes in protein levels (Fig. 3D and 4A). The expression of these proteins correlated in cortex with the MeCP2 protein levels and their steady-state level was restored by gene therapy (Fig. 4B). Quantitative immunoblot confirmed the steady-state level changes of most of these proteins after *Mecp2* loss of function (Fig. 4C-D). Furthermore, genes encoding these proteins showed significant peaks of MeCP2 binding activity near the annotated translation initiation site (Fig. 4E) (*30*). We then used Aacs, Calb1 and Etnppl to benchmark the temporal evolution of their transcript levels as well as the other members of the 119 cortico-hippocampal proteins after acute conditional loss of the Mecp2 gene (Fig. 4F). We found that the transcripts of Aacs, Calb1 and Etnppl and 26 other proteins significantly changed their level after one week of *Mecp2* gene ablation (Fig. 4F). Nearly one third of the 29 early responder mRNAs were metabolic annotated genes (Reactome pathway R-HSA-1430728, Odd ratio 4.5, corrected p=0.016. Grhpr, Nsdhl, Slc6a7, Fech, Auh, Hmgcs1, Adk, Bcat1, Aacs, Etnppl). The number of responder mRNAs significantly increased to 67 and 78 after 4 and 8 weeks of acute Mecp2 gene loss, respectively (Fig. 4F, p=1.8E-10). These results show that a fifth of the 119 *Mecp2*-sensitive proteins rapidly respond to modification of *Mecp2* gene expression likely corresponding to primary cellular mechanisms under MeCP2 control.

### Weighted Protein Co-expression Network Analysis of Brain *Mecp2* Loss- and Gain-of-Function Alleles and Their Rescue by Gene Therapy

We tested the robustness of q-defined findings with an orthogonal non-parametric analytical approach, **W**eighted **P**rotein **C**o-expression **N**etwork **A**nalysis, applied to all the quantified 10,433 proteins (Fig. 5, WPCNA, (*31*)). Our goal was to identify biomarkers at the protein level, therefore we analyzed the cortex and hippocampus proteomes to define clusters (modules) of highly correlated proteins. We then used module-specific first principal components to perform biweight midcorrelations (bicors) with MeCP2 protein content within and across brain regions and with the aggregated clinical score as traits (Fig. 1). We reasoned that top correlated modules with MeCP2 protein levels and aggregated clinical scores would 1) inform mechanisms affected by *Mecp2* deficiency, 2) identify *Mecp2*-sensitive proteins irrespective of whether they are directly under control of MeCP2 activity, and 3) inform putative biomarkers based on quantitative clinical traits. A total of 50 cortical and 44 hippocampal modules of co-expressed proteins were identified and ranked by their size, defined by each module constituent protein number, and module correlation with MeCP2 regional levels and aggregated clinical scores (Fig. 5A and 5S1A). Each module expression profile was expressed as an eigenprotein, corresponding to the first principal component of all protein expression profiles within that module (Fig. 5B and 5S1B) (*32*). Modules with similar expression patterns and trait correlations clustered near each other (Fig. 5B and 5S1B). We selected six modules in cortex and four in hippocampus that had the strongest correlation with MeCP2 protein levels and clinical traits (Fig. 5B and 5S1B arrows). More than 75% of all the protein constituents of each one of these modules correlated with MeCP2 protein levels and clinical score traits (Fig. 5S2, corrected p value<0.05). Eigenprotein profiles of the selected modules displayed coherence in their response to gene dosage and gene therapy, strongly correlated with MeCP2 protein levels, and with the aggregated clinical score (Fig. 5C and 5S1C). Moreover, these modules were enriched in synaptic, mitochondrial, central carbon metabolism, nuclear, and DNA repair pathways, corroborating ontologies identified by an orthogonal q-defined strategy (Fig. 5D, 5S1D, and 3S2). Our results establish a paradigm to define a robust collection of proteins and ontologies responsive to MeCP2 protein expression to inform the selection of putative Rett syndrome biomarkers in human samples. Our paradigm rests on a rigorous combination of large cohorts of *Mecp2* gain- and loss-of-function mutations, *MECP2* AAV9 gene therapy, multiomics, orthogonal analytical bioinformatic tools, MeCP2 protein levels, and animal clinical assessments to produce a robust mouse-defined database for human biomarker discovery selection.

**Figure 5.**
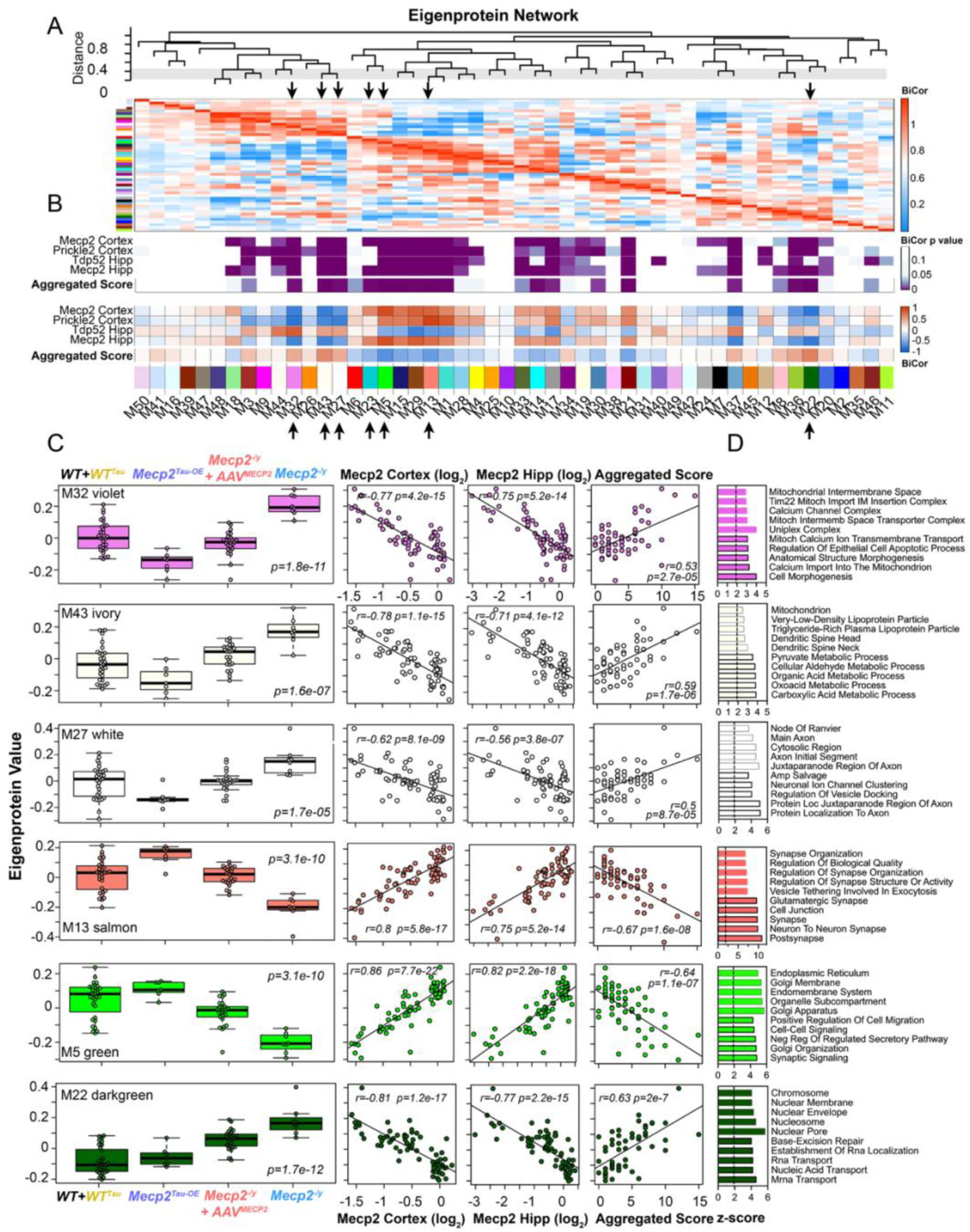
Cortical Protein Coexpression Network of *Mecp2* Mouse Models without and with MECP2 Gene Therapy. A. Weighted protein co-expression network analysis (WPCNA) grouped proteins into distinct protein modules (M1–M50) clustered to assess module relatedness based on correlation of protein co-expression eigenproteins. B. Biweight midcorrelation (BiCor) analysis of module eigenproteins with either Mecp2, Prickle2 and Tdp52 protein levels, and the aggregated behavioral score as traits. Purple heat map shows significance of association of each trait to a module using Benjamini-Hochberg FDR-corrected Student correlation p value. Red-blue heat map indicates strength of association where red represents positive correlation and blue indicates negative correlation of eigenproteins to each module. Module-trait correlations were performed using biweight midcorrelation (bicor). C. Changes in eigenprotein abundance in salient modules indicated by arrows in A and B across *Mecp2* mouse models without and with *MECP2* Gene Therapy. Multiple ANOVA with Tukey post hoc correction was used to assess the statistical significance of abundance changes. Eigenprotein correlation with indicated modules for each subject using Mecp2 content in two brain regions and the aggregated score as traits (Biweight midcorrelation). D. Selected modules with significant trait correlations were analyzed by gene ontology (GO) analysis, dotted line marks Z-score of 2. Processed data used to build figures can be found in supplementary table II.

### Proteomic Prediction of Brain Metabolic Defects in Mecp2 Loss-of-Function Mice

The 119 cortico-hippocampal *Mecp2*-responsive protein-mRNA pairs are enriched in intermediate metabolism annotated genes recapitulating prior *Mecp2-KO* proteomes (Fig. 3S2, 5D, and 5S1D). (*24, 27*) and a metabolomics study of patients with RTT (*33*). We reasoned that proteome-predicted metabolic alterations would serve as an independent, unbiased, and encompassing validation strategy. The 119 cortico-hippocampal *Mecp2*-responsive protein-mRNA pairs include aerobic respiration, central carbon, amino acid, and NAD^+^-NADH metabolism, such as the malate-aspartate shuttle (Fig. 3S2, 5D, 5S1D, and 6). In fact, these 119 protein-mRNA pairs are enriched 2.7-fold in Uniprot proteins annotated to nicotinamide adenine dinucleotide (NAD)-binding proteins such as Aldh6a1, Me2, and Sdha (p<0.004, Exact hypergeometric test, Fig. 6A, F, and G). We focused on proteins that participate (Slc25a13 and Got1, Figs, 6A-B) or are coupled (Bcat1) to the malate-aspartate shuttle, which is necessary for the transport of electrons from NADH across the inner mitochondrial membrane into the matrix (*34, 35*). These proteins were increased in *Mecp2-KO* brain, a phenotype rescued by MECP2 gene therapy. Moreover, their levels were reduced in Tau-*Mecp2* mouse brain (Fig. 6A-B).

**Figure 6.**
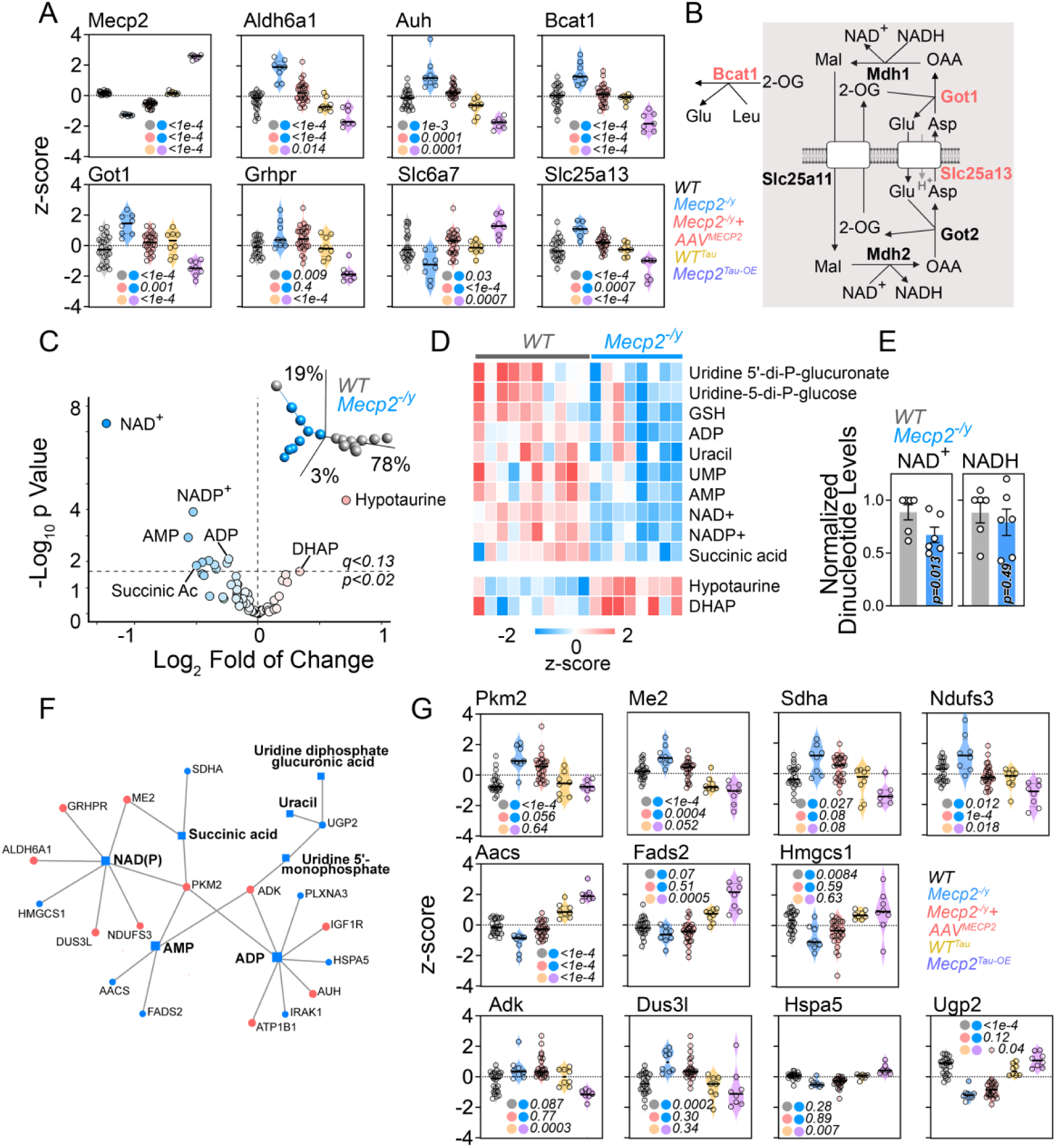
Metabolic Phenotypes in *Mecp2^-/y^* Cortex are Predicted by the *Mecp2*-sensitive 119 protein-mRNA pairs. A. Mass spectrometry levels of proteins annotated to metabolic ontologies. B Diagram of the malate-aspartate shuttle. Membrane depicts the inner mitochondria membrane. Red font marks proteins increased in *Mecp2*-KO brains and presented in panel A. C. Targeted metabolomics volcano plot comparing wild-type and *Mecp2*-KO cortices. Threshold q value (Benjamini-Hochberg FDR-corrected q<0.13, p<0.024, t-test) was chosen by the PCA and Euclidean distance discrimination of brain samples by genotype (shown in insert). D. Z-scored heat map of metabolites significantly changed in C. E. Enzyme-coupled determination of NAD^+^ and NADH in independent cohorts of wild-type and *Mecp2* KO cortices. Values were normalized to a control animal per cohort. Unpaired median difference two-sided permutation t-test, n=6. F. MetaboAnalyst network integration of metabolites in D and the 119 protein-mRNA pairs sensitive to *Mecp2* gene dosage. Red and blue symbols represent analytes increased or decreased significantly, respectively. G. Mass spectrometry levels of proteins integrated in the network in panel F. Panels A and G One-Way Anova followed by Benjamini, Krieger and Yekutieli multiple corrections. Processed data used to build figures can be found in supplementary table I.

To assess the metabolic consequences of the *Mecp2* proteome alterations, we performed a LCMS-based metabolomic study of the *Mecp2-KO* cortex focusing on intermediaries of central carbon, amino acid, and nucleotide metabolism (Fig. 6C-D). We identified 12 metabolites that discriminated genotypes by PCA, with NAD^+^, NADP^+^, and AMP as the most reduced metabolites in *Mecp2-KO* cortex (Fig. 6C-D). We confirmed decreased levels of NAD^+^ in *Mecp2-KO* cortices using an enzymatically-coupled assay to measure nicotinamide adenine dinucleotides (Fig. 6E). To infer protein-metabolite networks altered in *Mecp2*-KO cortex, we fed the 119 proteins and 12 metabolites sensitive to *Mecp2* into the MetaboAnalyst 6.0 tool (Fig. 6F) (*36*). The protein-metabolite network revealed proteins whose levels in *Mecp2*-KO were restored by gene therapy (Me2, Ndufs3, Aacs; Fig. 6G) and proteins whose levels were sensitive to gene dosage (Ndufs3, Aacs, Fads2, Adk, Hspa5, Ugp2; Fig. 6G). These results provide independent validation of the proteome-transcriptome alterations in *Mecp2*-KO brain and confirm prior findings of central carbon metabolism defects in *Mecp2* deficiency (*24, 27*).

### Identification of Differentially Abundant CSF Proteins in Rett Syndrome by NULISA Multiplex ELISA

To assess if a mouse-defined *Mecp2*-sensitive protein database could inform putative human biomarker selection, we independently analyzed cerebrospinal fluid protein composition of 13 neurotypical children and 16 patients with sequence confirmed RTT from two geographically distinct cohorts (Fig. 7A) (*37, 38*). Subjects with RTT were sex-matched to neurotypical children, who were slightly older (median age 8 and 12 years, respectively) (Fig. 7B). We used the highly sensitive NUcleic acid Linked Immuno-Sandwich Assay (NULISA) to measure 120 CNS-curated proteins, representing neurodegenerative disease-related protein markers and inflammation and immune response-related cytokines and chemokines, per sample by coupling oligo-tagged antibody-pairs with next-generation sequencing quantitative readout (*39*). The performance of this platform has been cross-validated with clinically established biomarkers of Alzheimer’s disease in CSF and plasma (*40–42*). There were no differences in the levels of proteins indicative of blood contamination, such as hemoglobin HBA1, found in control and RTT CSF (Fig. 7S1). We used two analytical tools to identify proteins whose levels were altered in individuals with RTT. First, we processed NULISA data by discriminant partial least squares regression analysis to reduce multivariate datasets from the 120 variables to two latent variables (LV1 and LV2) (Figs. 7C). Latent variables allowed us to identify differences between neurotypical children with those with RTT from unrelated noise in measurements (*43, 44*). Each LV represents a profile of proteins that vary between neurotypical and RTT patients. The first latent variable (LV1) captured the major source of variation associated with RTT status, separating neurotypical children from RTT patients irrespective of geographical location, while LV2 described variation unrelated to diagnostic group (Fig. 7C). We used a variable importance in projection (VIP score) for each protein identified in LV1 to determine a protein’s discriminatory capacity between neurotypical and RTT groups within the PLS-DA model (Fig. 7D). This approach identified 52 proteins capable of discriminating between neurotypical and RTT groups (Fig. 7D). We combined this strategy with volcano-plot selection which identified 43 of the 52 proteins as changing in RTT patient samples (Fig. 7E-F). To designate a protein as a putative RTT-affected protein, we used as criteria the intersection of the PLS-DA-VIP and volcano plot selection NULISA datasets either between themselves, with the q<0.001 *Mecp2-KO* cortical proteome, or the q-defined 119 *Mecp2*-sensitive protein-mRNA pairs from cortex and hippocampus (Fig. 7A). Mouse orthologues of PTN and IGF1R were altered in *Mecp2-KO* mouse cortex and hippocampus at the mRNA and protein levels. In contrast, mouse orthologs of VGF, ANXA5 (WPCNA module M27 0.77), SNAP25 (M1 0.86), NPTX1, NPTX2, and NPTXR were affected in *Mecp2-KO* mouse cortex only as proteins but not as mRNAs (Fig. 7G-H).

**Figure 7.**
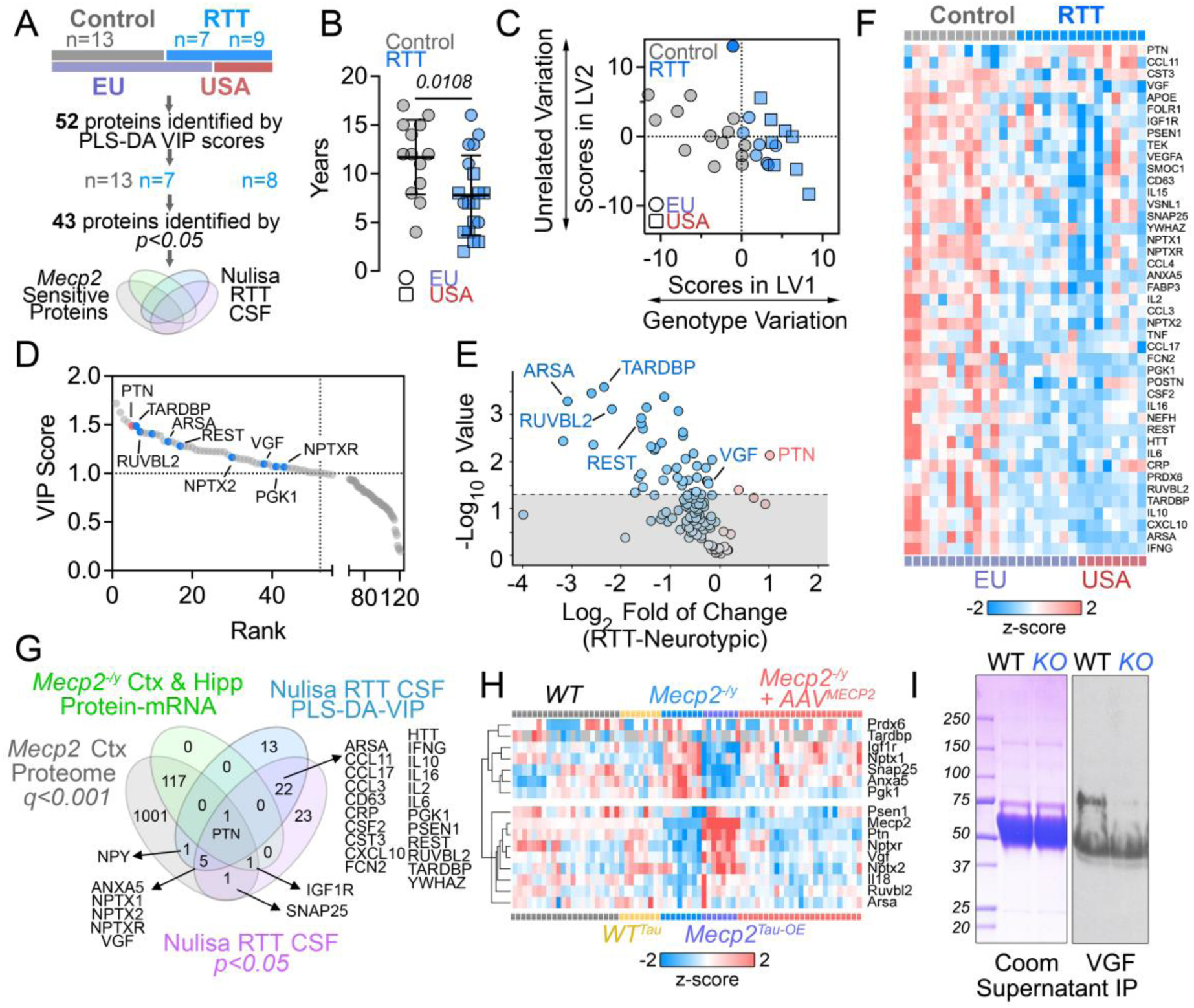
Composition of Neurotypical and Rett Syndrome Subjects Cerebrospinal Fluid Measured by NUcleic acid Linked Immuno-Sandwich Assay (NULISA). A. Neurotypical and Rett syndrome cohort’s characteristic and experimental design. B. Female subjects age by genotype. Unpaired t-test (Average±SD). Symbol shapes depict cohorts by geographic location (USA, United States; EU, European Union). C. Discriminant partial least squares regression model constructed from the NULISA dataset regressed by genotype. The model identifies a latent variable (LV1) that scores based on NULISA protein expression measurements and predicts diagnosis. LV2 describes variation that is not connected to diagnosis. D. VIP score >1 identifies 52 proteins with discriminatory capacity between neurotypical and RTT subjects within the PLS-DA model. Rank presents all the proteins measured with the NULISA platform ordered by VIP score. E. Volcano plot of NULISA samples comparing RTT and Neurotypical CSF. t-test p<0.05 (Benjamini-Hochberg FDR-corrected q<0.13). One RTT subject sample detected as outlier in the LV2 dimension (dark blue symbol in C) was excluded. F. Z-scored heat map off 43 proteins selected in E. Red denotes increased protein levels in RTT CSF. G. Venn diagram outcomes of overlapping analytes using selection process shown in A. H. Z-scored heat map off proteins selected in G show coherent differential expression profiles in *Mecp2* gain- and loss-of function mutations and their restoration by AAV9-RTT271 therapy. H. *MECP2*-null SH-SY5Y neuroblastoma-conditioned media show decreased levels of VGF. Processed data used to build figures can be found in supplementary table I.\

We focused on human VGF for three reasons. First, mouse Vgf belongs to the M13 module (bicor to M13: 0.63), one of the strongest cortical modules (Fig. 5). Second, Vgf is a secreted protein present in large-dense core vesicles in neurons (*45*). Third, we reported that Vgf levels are reduced in cerebrospinal fluid of either *Mecp2^-/y^* and *Mecp2^-/+^* mice (*27*). Expression of mouse Vgf correlated with Ptn and MeCP2 protein levels but not with other growth factors including Bdnf (Fig. 7S2) (*46*). We confirmed the sensitivity of Vgf to *Mecp2* in an independent system by CRISPR KO of the *MECP2* gene in human SH-SY5Y neuroblastoma cells (Fig. 7I) (*24*). VGF content was decreased in these *MECP2-KO* cells both in conditioned media and cell extracts (Fig. 7I and 7S3). These results show that 28 out of 120 proteins measured by NULISA (23%) are altered in our RTT patient cerebrospinal fluid samples. Of these 28 proteins, 9, or 7.5% of the NULISA-measured analytes, are proteins curated by their sensitivity to *Mecp2* gene dosage and/or gene therapy in mouse models of the disease.

### Murine Proteome–Trait Correlations and Discriminatory Capacity of Mouse-Informed Putative Biomarkers in Rett Syndrome

Molecules selected as putative disease biomarkers should quantitatively respond to the content/activity of the causative gene and reflect subject well-being, function, and/or survival. We tested if proteins selected as putative biomarkers from direct/unbiased analyses of CSF samples from patients with RTT predicted mouse brain MeCP2 protein levels and animal clinical status (Fig. 8). We used the aggregated clinical score as a trait reporting neurological and animal well-being and function and the whole cortical proteome dataset to extract a linear metric correlating protein levels and clinical outcomes. We calculated Beta (effect size) corresponding to the slope of a fit line for each protein as a predictor of either MeCP2 protein content or the aggregated clinical score as outcomes as well as an effect size FDR-corrected p value (Fig. 8A-B). We calculated Beta for proteins in three datasets: the 119 protein-mRNA pairs responsive to *Mecp2* dosage in cortex and hippocampus, proteins identified in *Mecp2-KO* mouse CSF (*27*), and the proteins selected from the Rett CSF NULISA datasets. These three datasets clustered in the same cortical modules defined by WPCNA (Fig. 8S1) and partially overlapped (Fig. 8C). Among the proteins with the strongest effect size were Pgk1 and Ruvbl2 (Fig. 8A, B and D). An increase of 1 standard deviation (SD) in Pgk1 predicts a decrease of ∼ 7SD in MeCP2 and a worsening of ∼40 SD in the clinical score. Conversely, an increase of Ruvbl2 by 1SD predicts an increase of 8 SD in MeCP2 levels and an improvement of 50 SD in clinical outcome. Vgf provides an intermediate predictive level where each SD in Vgf content increase predicts increases of 3 SD in MeCP2 abundance and an improvement of 16 SD in the clinical score. Molecules with large effect size in mouse datasets were able to discriminate neurotypical and Rett CSF profiles as determined by the area under the curve (AUC) in Receiver Operating Characteristic (ROC) analysis (Fig. 8E and 8S2). We conclude that selection of putative disease biomarkers in human samples can be informed by genetically curated mouse datasets and animal clinical outcomes. We propose that this strategy can be utilized in either a genetic model-to-human or a human-to-genetic model direction and applied to identify fluid biomarkers and mechanisms in genetic diseases such as those affecting neurodevelopment.

**Figure 8.**
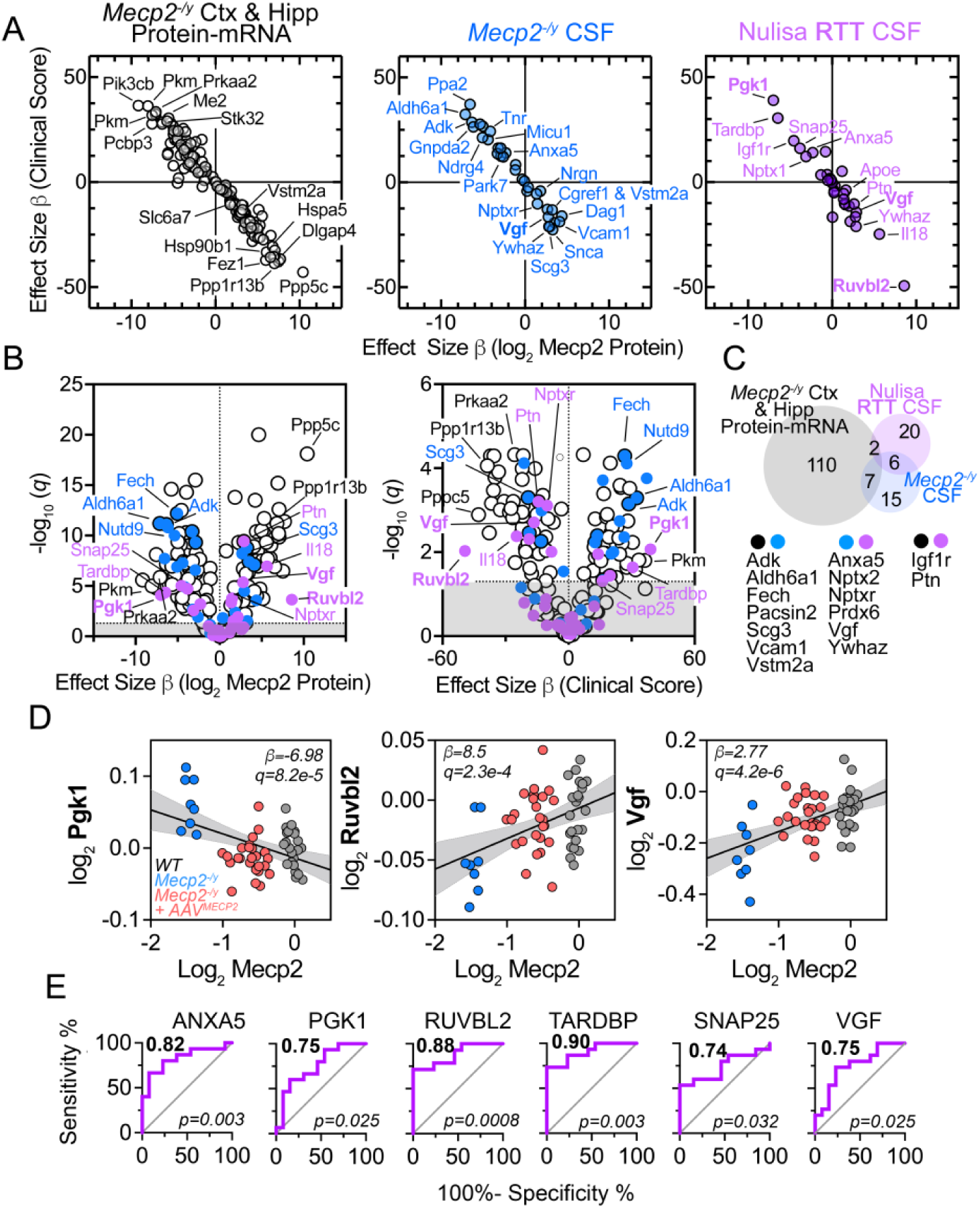
Effect Size and Discriminant Power of Proteins Curated by *Mecp2* Genotypes and MECP2 Gene Therapy on RTT Diagnosis. A. Beta (effect size) corresponding to the slope of a fit line for each protein as a predictor of either MeCP2 protein content or the aggregated clinical score as traits. Three datasets are analyzed: 119 proteins responsive to *Mecp2* dosage in cortex and hippocampus (open symbols), proteins identified in *Mecp2-KO* mouse CSF (blue symbols), and the proteins selected from the RTT CSF NULISA datasets (purple symbols). B. Volcano plots of trait effect size Beta vs −log10 FDR corrected p value for the correlation. Symbol colors as in A. C. Venn diagram of protein overlaps of the three datasets in A and B. D. Examples of three proteins selected from the Nulisa RTT CSF plot in B depicted as a correlation with MeCP2 protein levels. Color annotates genotype and gene therapy. E. Discrimination between neurotypical and RTT CSF by selected analytes determined by the area under the curve (AUC, bold number upper left corner) in Receiver Operating Characteristic (ROC) analysis. Processed data used to build figures can be found in supplementary table II.

## Discussion

We present a genetically guided, unbiased multiomics strategy to identify putative mechanisms and candidate biomarkers of MeCP2 dysfunction across species. We defined a broad *Mecp2*-sensitive proteome, validated via multiple approaches. First, the breadth of this proteome is supported by the study of large animal cohorts and stringent statistical filtering—rather than fold-change thresholds—to account for the cellular heterogeneity of brain tissue and the modest transcriptomic effects observed in *Mecp2* knockout models by others and us (*6, 7, 9, 15, 24, 47*). Second, we prioritized proteins that change in a coherent manner in both loss- and gain-of-function *Mecp2* alleles and confirmed them at the transcript level with published datasets of loss- and gain-of-function models (*6, 9, 11, 25, 29*). Our approach may exclude molecules uniquely affected by *Mecp2*-null or *Mecp2*-overexpression mutations, but it emphasizes a subset of gene products where *Mecp2* dosage acts as a rheostat, coherently regulating expression either directly or downstream of direct Mecp2 targets. For example, Vgf appears to be an indirect downstream target, as its protein levels respond to *Mecp2* dosage and gene therapy despite stable mRNA expression. In contrast, Ptn behaves as a direct target of *Mecp2* where both protein and mRNA respond coherently to *Mecp2* gene dosage across multiple independent analyses in mouse and human. Ptn is a secreted neurotrophic factor implicated in neuronal development, synaptic plasticity, and neuroprotection, making its dysregulation a possible mediator and reporter of MeCP2 dysfuncion (*48–50*). Future studies using ubiquitously expressed gain-of-function alleles, that model MECP2 duplication syndrome (*51*), may reveal a broader *Mecp2*-sensitive proteome across additional cell types, as our choice of the Tau-*Mecp2* overexpression model introduces bias toward neuronal-expressed genes (*22*). We further validated our proteome by biochemical testing of selected proteins (Aacs, Calb1, Got1, Vgf, Ptn) and metabolic pathways, using an orthogonal targeted metabolomics approach. Most importantly, we demonstrate that *MECP2* gene therapy rescues this proteome and restores clinical phenotypes in mice, thus supporting the notion that the proteome and clinical phenotypes are causally linked. Gene replacement therapy was achieved using an AAV9 construct engineered for precise control of *MECP2* expression that is currently under evaluation in human trials (*3, 19*). We conclude that the rescue of a *Mecp2*-sensitive proteome by gene therapy reveals the breadth and complexity of molecular disruptions in preclinical models of RTT, providing opportunities for early therapeutic interventions. Moreover, the identified *Mecp2*-sensitive proteins orient us to yet to be explored RTT’s mechanisms of disease and potential biomarkers of brain MeCP2 dysfunction in affected individuals.

The *Mecp2*-sensitive proteome highlights several previously reported processes disrupted by *Mecp2* mutations in mouse models and individuals with RTT, including synaptic transmission and organization, cholesterol metabolism, and mitochondrial central carbon metabolism—particularly monocarboxylate metabolism, which is closely linked to mitochondrial amino acid metabolism (*12, 24, 33, 52*). One such example is the malate-aspartate shuttle matrix (*34, 35*). Given that many of these metabolic pathways require NAD⁺/NADH as cofactors, we performed targeted metabolomic analyses as an all-encompassing strategy to test proteome-derived hypotheses. We found reduced NAD⁺ levels in *Mecp2*-deficient brains. The exact cause of this reduction remains unknown but is likely multifactorial, as multiple gene products in the *Mecp2*-sensitive proteome require NAD as cofactors. Regardless, NAD⁺ and NADH levels may serve as rapid-response biomarkers of disease, assessable via non-invasive methods such as magnetic resonance spectroscopy of human brains (*53, 54*). NAD may also link several seemingly unrelated mechanisms that genetically interact with *Mecp2*-null mutations to influence survival. These include cholesterol metabolism, ketone body metabolism, and DNA repair (*55*). Acetoacetyl-CoA synthetase (Aacs) does not directly use NAD⁺/NADH as cofactors but intersects with pathways that do (*56*). Aacs is among the gene products that responded coherently to both *Mecp2* loss- and gain-of-function alleles as well as to gene therapy at the level of protein and mRNA in our study. This suggests that Aacs is a direct target of Mecp2, an idea supported by the binding of MeCP2 to the *Aacs* genomic locus, and the rapid Aacs mRNA response to acute deletion of the *Mecp2* gene. Given that mutations in *Aacs* can rescue the *Mecp2*-KO survival phenotype (*57*), this supports the idea that our *Mecp2*-sensitive proteome provides a rich source for identifying disease-modifying mechanisms.

A surprising finding is the lack of coherence in the non-coding transcriptomes from cortex and hippocampus and the coding transcriptome in hippocampus. We do not attribute this to technical issues, as the same animals processed in parallel displayed a coherent coding transcriptome in the cortex. Our data analysis approach captures non-coding transcriptome changes that seem to be driven by the strain of mouse rather than the *Mecp2* gene dosage and gene therapy. This suggests that true *Mecp2*-sensitive RNA expression changes are subtle compared to the effect of the animal strain in the non-coherent transcriptomes, thus evading their discovery in our experimental design. Published cortical coding transcriptomes reveal regional and cell-type-specific differences in mRNAs affected by *Mecp2* knockout (*7, 11, 24*). Our study confirms this finding, consistent with previous reports by our group and others. Among the 1,852 *Mecp2*-sensitive proteins identified in the cortex and hippocampus, only 327 were shared between the two regions. Of these shared proteins, only 119 had corresponding mRNAs that exhibited parallel expression changes across genotypes and following gene therapy. This limited overlap between transcriptomic and proteomic changes was similarly observed in both brain regions: only 12% of differentially expressed mRNAs were associated with protein-level changes consistent with *Mecp2* gene dosage and rescue. We previously hypothesized that this discordance reflects post-transcriptional regulatory mechanisms influenced by Mecp2 (*24*). Although we examined various small non-coding RNAs as potential mediators, none met the genetic coherence criteria used to define our *Mecp2*-sensitive proteome. Alternatively, alterations in protein synthesis or degradation may underlie this discrepancy (*58*). We propose that discordance between the transcriptome and proteome may constitute a molecular phenotype of *Mecp2* deficiency in the brain.

The concurrent collection of clinical and proteomic data from individual mice, both with and without *MECP2* gene therapy, enabled us to evaluate how *Mecp2*-sensitive protein levels correlate with MeCP2 protein expression and clinical outcomes in mice. The analyses demonstrate that these proteins can serve as surrogate markers of disease in murine disease models. Our goal was to leverage these *Mecp2*-sensitive brain proteins to guide the identification of candidate biomarkers of MeCP2 dysfunction in patients with Rett syndrome. We focused on cerebrospinal fluid, as we had previously observed proteomic changes in the cerebrospinal fluid of *Mecp2^-/y^* and *Mecp2^-/+^* mice (*27*), and because it may helpful in monitoring genetic therapies currently under development. Furthermore, biomarkers in cerebrospinal fluid are brain specific and have the potential to be detected in extracellular vesicles in blood (*59*). We analyzed two cohorts of patients with RTT and matched neurotypical controls using a 120-plex ELISA panel targeting proteins implicated in various neurological disorders. This analysis identified 43 proteins that differentiated RTT from control cerebrospinal fluid, 9 of which were present in the *Mecp2*-sensitive proteome, suggesting they are good candidates to monitor MeCP2 dysfunction in disease and therapeutic interventions. These proteins exhibited robust effect sizes, supporting their potential as individual or composite surrogate biomarkers. While these findings are encouraging, further validation is needed. The limited human sample size restricted our ability to perform multivariate analyses correlating biomarker levels with clinical severity or mutation type. Independent replication and prospective studies are required to substantiate these findings.

We used a multiplexed NULISA to test the translational potential of the *Mecp2*-sensitive proteome in human cerebrospinal fluid. We selected the NULISA platform for its sensitivity, large dynamic range, and low sample volume requirements (*39*). This technology has recently demonstrated clinical utility in Alzheimer’s disease plasma studies, performing comparably to or exceeding existing diagnostic tools such as Simoa assays and Tau-PET imaging (*40, 41*). These features enhance the potential for adapting our cerebrospinal fluid findings to plasma-based biomarker discovery in RTT. A major advance in RTT therapy is the initiation of genetic therapy, specifically gene replacement trials. Notably, one such trial employs an EXACT AAV9 construct, similar to that used in our mouse studies (*19*). Plasma samples from these patients, collected before and after treatment, would represent a powerful opportunity to test the utility of *Mecp2*-sensitive proteins as circulating biomarkers of disease progression and therapeutic response.

We conclude that the selection of putative disease biomarkers in human samples can be effectively guided by genetically curated datasets and clinical outcomes from animal models. This strategy can be applied bidirectionally—either from genetic models to human studies or from human findings back to genetic models—and offers a broadly applicable framework for identifying biomarkers and underlying mechanisms across a wide range of genetic disorders.

### Limitations

Several limitations of this study should be acknowledged. First, our discovery platform relied on hemizygous male *Mecp2*-null mice rather than female heterozygous animals, a design chosen to maximize statistical power and enable robust therapeutic rescue studies but one that does not fully recapitulate the mosaic biology of Rett syndrome. Second, the proteins identified should be considered biologically prioritized putative biomarkers rather than qualified pharmacodynamic biomarkers. Although many candidates showed concordant changes across transcriptomic, proteomic, gene therapy, and human datasets, their temporal responsiveness, dynamic range, and predictive value following acute changes in MECP2 dosage remain to be established through longitudinal and dose-escalation studies. Third, our gene therapy experiments used vehicle-treated rather than AAV-control animals, although additional analyses found no evidence of vector-associated inflammatory or toxicity signatures. Finally, technical limitations associated with mouse CSF volume precluded parallel CSF proteomic analyses. Future studies in female models, independent patient cohorts, and longitudinal therapeutic paradigms will be important to further validate the translational utility of these candidate biomarkers.

## Materials and Methods

### Mice and Gene Therapy

Six-week old, C57BL/6J and *Mecp2^tm1.1Bird^* (The Jackson Laboratory #000664 and #003890) mice were euthanized and tissue collected as approved by the Emory University Institutional Animal Care and Use Committees. Briefly brains were sliced coronally and the cortex and hippocampus from left and right hemispheres of adjacent slices in were flash frozen (*60*).

For the gene therapy experiments, animal handling, detailed procedures, and scoring have been described in (*61*). All animal procedures adhered to European (86/609/EEC) and UK (Animals [Scientific Procedures] Act 1986) ethical guidelines. The B6.129P2(C)-Mecp2tm1.1Bird/J mouse model of Rett syndrome (RTT) was maintained on a mixed C57BL/6 CBA background. Heterozygous Mecp2+/− females were crossed with WT B6CBAF1 males to generate experimental cohorts through timed matings. Pregnancy was confirmed at gestational day 15–17, and pregnant females were then housed separately and monitored daily for births. Neonates were genotyped at postnatal day 0 (P0) and ear-notched for identification at P14–21. Mice were kept under controlled environmental conditions: 12-hour light/dark cycles, temperature of 20° ± 2°C, and humidity of 50 ± 15%, with HEPA-filtered air exchanges (20/hour). Water and food were provided ad libitum. Similar husbandry was used to keep Mecp2Tau-OE (*22*)

For in vivo gene transfer studies, mice were dosed intracerebroventricularly (ICV) at P0–2. Neonates were anesthetized with isoflurane and injected bilaterally with either AAV9-RTT271 vector or vehicle (PBS + poloxamer) using a 30-gauge dental needle attached to a Hamilton syringe. Injections targeted the temporal cortex (∼1–2 mm lateral to midline and ∼2 mm anterior to lambda) at a depth of ∼2–3 mm. Post-injection, pups recovered on heated pads before being returned to their home cage.

The study evaluated RTT-like phenotypes from 4 weeks of age onward. A RTT scoring system was employed to assess disease severity across six domains: mobility, gait, hindlimb clasping, tremor, breathing, and general condition (*23*). Each parameter was scored from 0 (normal) to 2 (severe), yielding a cumulative score of 0–12. A single evaluator was blinded to treatment groups throughout the study. Open field and whole-body plethysmography were conducted at the age of 8 weeks (*62*). Mice were euthanized using terminal dose of anesthesia and the brains were collected and sliced as mentioned above.

### RNA sequencing and other analyses

Admera Health performed all RNA isolation, library preparation, and sequencing. For both the coding RNA experiments and small RNA experiments, RNA was isolated from the Hippocampus or Cortex using Qiazol phase separation, followed by cleanup with RNeasy 96. Isolated RNA sample quality was assessed by both RNA Tapestation assay and High Sensitivity RNA Tapestation assay (Agilent Technologies Inc., California, USA) and quantified by Infinite F Nano+ 200 Pro Tecan (Tecan, Switzerland).

Coding RNA transcripts were isolated using NEBNext® Poly(A) mRNA Magnetic Isolation beads as part of the NEBNext® Ultra™ II Directional RNA Library Prep Kit for Illumina® (New England BioLabs Inc., Massachusetts, USA). This kit was also used for cDNA synthesis and library preparation. Prior to first strand synthesis, samples are randomly primed (5’ d(N6) 3’[N=A,C,G,T]) and fragmented. The first strand of cDNA was synthesized with the Protoscript II Reverse Transcriptase with a longer extension period, approximately 30 minutes at 42⁰C. Final libraries quantity was assessed by Qubit 2.0 (ThermoFisher, Massachusetts, USA) and quality was assessed by TapeStation HSD1000 ScreenTape (Agilent Technologies Inc., California, USA). Final library size was about 450bp with an insert size of about 300bp. Illumina® 8-nt dual-indices were used. Equimolar pooling of libraries was performed based on QC values and sequenced on an Illumina NovaSeq X Plus (Illumina, California, USA) with a read length configuration of 150 PE for 60M PE reads per sample (30M in each direction).

The NEB NEBNext Small RNA Library Preparation Kit for Illumina (New England BioLabs Inc., Massachusetts, USA) was used for the small RNA libraries. The final library quantity was assessed by Qubit 2.0 (ThermoFisher, Massachusetts, USA), and quality was evaluated by TapeStation HSD1000 ScreenTape (Agilent Technologies Inc., California, USA). The final library size was about 250bp with an insert size of about 100bp. Illumina 6-nt single-indices were used. Equimolar pooling of libraries was performed based on QC values and sequenced on an Illumina NovaSeq X Plus (Illumina, California, USA) with a read length configuration of 150 PE for 20M PE reads per sample (10M in each direction).

For RNA sequencing analysis:FastQC was used to remove samples of poor quality (*63*). Adapter sequences and readings of poor quality were trimmed using Trimmomatic (version v0.39). We then used the web interface and public servers, usegalaxy.org and usegalaxy.eu, where sequence reads were uploaded for analysis (*64*). The Galaxy server running Hisat2 (Galaxy Version 2.2.1+galaxy0), FeatureCounts (Galaxy Version 1.6.4), and Deseq2 (Galaxy Version 2.11.40.8+galaxy1) was used to map sequence reads (*65–67*). FeatureCounts files and raw files are available at GEO with accession GSE300534. We utilized Hisat2 with the following settings: paired-end, stranded, default settings (except for when a GTF file was used for transcript assembly). For GTF files, we used the Mus musculus (Mouse), Ensembl, GRCm39 build from iGenome (Illumina). The aligned SAM/BAM files were processed using Featurecounts with Default settings, except we used the Ensembl GRCm39 GTF file and output for DESeq2 and a gene length file.

Small RNAs were analyzed within Illumina’s Basespace platform. Raw samples were trimmed using Flexbar to remove the adapter sequence (*68*). FastQC was used to remove samples of poor quality. Mapping was done using the mm10 reference genome. Alignments and counts were done using the STAR aligner (v. 2.0.2) as it outperforms Hisat2 for small RNA sequencing analyses (*69–71*). MultiQC (v. 1.19) was used to visualize alignment. The following databases in parentheses were used to compile each category of small RNAs: circRNA (circBase), miRNA (miRbase), piRNA (piRNABank), snoRNA (gencode), snRNA (gencode), & tRNA (gtRNAdb). FeatureCounts was used to quantify counts and DESeq2 (using the Galaxy server) was used to determine differential expression. Raw files, Individual counts for each small RNA and normalized counts for the small RNAs are available at GEO with the Accession number GSE300536.

Gene counts were normalized using DESeq2 (*67*) followed by a regularized log transformation. Differential Expression was determined by DESeq2 using the following settings: For Tau-MECP2 overexpression and RTT271 gene therapy experiments (both RNAseq and small RNA analysis) the factors were tissue type (hippocampus and cortex), Injection type (RTT271_1e11 and RTT271_3e11), MeCP2 status and Tau-MECP2 overexpression. Pairwise comparisons were done across injection type, MeCP2, and Tau-MECP2 overexpression status. All normalized tables, including regularized log transformation and variance stabilized tables, were generated using size estimation, where the standard median ratio was used; the fit type was parametric, and outliers were filtered using a Cook’s distance cutoff.

RTT271_Counts and Mecp2_E2_CDS_Tau_Counts were determined in appropriate samples. RTT271 FASTA sequence was aligned to each injected sample using Bowtie2 (Galaxy version v2.5.3+galaxy0), and aligned reads are reported. For Tau-Mecp2 overexpression, the Mecp2_E2_CDS_Tau FASTA sequence was aligned to the Tau-Mecp2 overexpression samples using Bowtie2, and the number of reads aligning is reported.

Cut&Run data analysis was implemented with Alignment and Counts collected as described by Bajikar et al. (2023) (*30*). Sequences and Normalized Bigwig files were downloaded from GEO with accession code GSE213752. Alignment of raw FASTQ files were aligned using bowtie2 (Galaxy Version 2.5.5+galaxy0) in Galaxy with the following settings –dovetail. Alignment was performed using Galaxy built in genome mm10 and E.coli K12 Genome (GCF_000005845.2_ASM584v2). Bedtools MultiCovBed (Galaxy Version 2.31.1) was used to count fragments over a 7kB segment centered around peaks in the 20kb region upstream from each gene. Integrative Genomics Viewer (IGV) v2.19.7 was used to visualize the average normalized Bigwig files centered on the gene of interests (Robinson et al., 2011). We performed differential expression of fragments in the 7kb peaks identified in the 20kb region for genes of interest using the DESeq2 package ((Galaxy Version 2.11.40.8+galaxy3)) using the E-coli spike in control counts as normalization factor and cooksCutoff set to FALSE.

### Proteomics

Tissue Homogenization and Protein Digestion. Samples were homogenized in 8 M urea lysis buffer (8 M urea, 10 mM Tris, 100 mM NaH2PO4, pH 8.5) with HALT protease and phosphatase inhibitor cocktail (ThermoFisher) using a Bullet Blender (NextAdvance). The lysates were sonicated for 2 cycles consisting of 5 s of active sonication at 30% amplitude, followed by 15 s on ice. Samples were then centrifuged for 10 min at 8,000 rpm and the supernatant transferred to a new tube. Protein concentration was determined by bicinchoninic acid (BCA) assay (Pierce).

For protein digestion, 70 μg of each sample was aliquoted and volumes normalized with additional lysis buffer. Samples were reduced with 5 mM dithiothreitol (DTT) at room temperature for 30 min, followed by 10 mM iodoacetamide (IAA) alkylation in the dark for another 30 min. Lysyl endopeptidase (Wako) at 1:20 (w/w) was added, and digestion allowed to proceed overnight. Samples were then 7-fold diluted with 50 mM ammonium bicarbonate. Trypsin (Promega) was then added at 1:2 (w/w) and digestion proceeded overnight. The peptide solutions were acidified to a final concentration of 1% (vol/vol) formic acid (FA) and 0.1% (vol/vol) trifluoroacetic acid (TFA) and desalted with a 10 mg HLB column (Oasis). Each HLB column was first rinsed with 1 mL of methanol, washed with 1 mL 50% (vol/vol) acetonitrile (ACN), and equilibrated with 2×1 mL 0.1% (vol/vol) TFA. The samples were then loaded onto the column and washed with 2×1 mL 0.1% (vol/vol) TFA. Elution was performed with 2 volumes of 0.5 mL 50% (vol/vol) ACN. A 60 ug equivalent aliquot was taken out and the residual combined to create a region specific global internal standard. All aliquots were dried down by speedvac.

Isobaric Tandem Mass Tag (TMT) Peptide Labeling. Each sample was re-suspended in 100 mM TEAB buffer (25 μL). The TMT labeling reagents (5mg) were equilibrated to room temperature, and anhydrous ACN (200μL) was added to each reagent channel. Each channel was gently vortexed for 5 min, and then 5 μL from each TMT channel was transferred to the peptide solutions and allowed to incubate for 1 h at room temperature. The reaction was quenched with 5% (vol/vol) hydroxylamine (2 μl) (Pierce). All channels were then combined and dried by SpeedVac (LabConco) to approximately 100 μL and diluted with 1 mL of 0.1% (vol/vol) TFA, then acidified to a final concentration of 1% (vol/vol) FA and 0.1% (vol/vol) TFA. Labeled peptides were desalted with a 30 mg HLB column (Oasis). Each HLB column was first rinsed with 1 mL of methanol, washed with 1 mL 50% (vol/vol) acetonitrile (ACN), and equilibrated with 2×1 mL 0.1% (vol/vol) TFA. The samples were then loaded onto the column and washed with 2×1 mL 0.1% (vol/vol) TFA. Elution was performed with 2 volumes of 0.6 mL 50% (vol/vol) ACN. The eluates were dried to completeness using a SpeedVac.

High-pH Off-line Fractionation. Dried samples were re-suspended in high pH loading buffer (0.07% vol/vol NH4OH, 0.045% vol/vol FA, 2% vol/vol ACN) and loaded onto a Water’s BEH 1.7 um 2.1mm by 150mm. An Thermo Vanquish was used to carry out the fractionation. Solvent A consisted of 0.0175% (vol/vol) NH4OH, 0.01125% (vol/vol) FA, and 2% (vol/vol) ACN; solvent B consisted of 0.0175% (vol/vol) NH4OH, 0.01125% (vol/vol) FA, and 90% (vol/vol) ACN. The sample elution was performed over a 25 min gradient with a flow rate of 0.6 mL/min. A total of 192 individual equal volume fractions were collected across the gradient and subsequently pooled by concatenation into 96 fractions and dried to completeness using a SpeedVac.

Liquid Chromatography Tandem Mass Spectrometry Cortex. All samples (∼1ug for each fraction) were loaded and eluted using Vanquish Neo (Thermofisher Scientific) an in-house packed 20 cm, 75um i.d. capillary column with 1.9 μm Reprosil-Pur C18 beads (Dr. Maisch, Ammerbuch, Germany) using a 34 min gradient. Mass spectrometry was performed with a Orbitrap Q-Exactive HFX (Thermo) in positive ion mode using data-dependent acquisition with 20 TopN cycles. Each cycle consisted of one full MS scan followed by as many as 20 MS/MS events. MS scans were collected at a resolution of 120,000 (410-1600 m/z range, 3×10^6 AGC, 50 ms maximum ion injection time). All higher energy collision-induced dissociation (HCD) MS/MS spectra were acquired at a resolution of 45,000 (0.7 m/z isolation width, 32% collision energy, 1×10^5 AGC, 96 ms maximum ion time). Dynamic exclusion was set to exclude previously sequenced peaks for 20 seconds within a 10-ppm isolation window.

Liquid Chromatography Tandem Mass Spectrometry Hippocampus. All samples (∼1ug for each fraction) were loaded and eluted using Dionex Ultimate 3000 RSLCnano (Thermofisher Scientific) an in-house packed 15 cm, 150 μm i.d. capillary column with 1.9 μm Reprosil-Pur C18 beads (Dr. Maisch, Ammerbuch, Germany) using a 25 min gradient. Mass spectrometry was performed with a high-field asymmetric waveform ion mobility spectrometry (FAIMS) Pro equipped Orbitrap Eclipse (Thermo) in positive ion mode using data-dependent acquisition with 1.5 second top speed cycles. Each cycle consisted of one full MS scan followed by as many MS/MS events that could fit within the given 1.5 second cycle time limit. MS scans were collected at a resolution of 120,000 (410-1600 m/z range, 4×10^5 AGC, 50 ms maximum ion injection time, FAIMS compensation voltages of −45 and −65). All higher energy collision-induced dissociation (HCD) MS/MS spectra were acquired at a resolution of 30,000 (0.7 m/z isolation width, 38% collision energy, 200% AGC, 54 ms maximum ion time, TurboTMT on). Dynamic exclusion was set to exclude previously sequenced peaks for 20 seconds within a 10-ppm isolation window.

Data Processing Protocol. All raw files were searched using Thermo’s Proteome Discoverer suite (version 3.0.1.27) with Sequest HT. The spectra were searched against a mouse uniprot database downloaded August 2020 (91414 target sequences). Search parameters included 10ppm precursor mass window, 0.05 Da product mass window, dynamic modifications methione (+15.995 Da), histidine, serine and threonine TMT (+304.207 Da), and static modifications for carbamidomethyl cysteines(+57.021 Da) and N-terminal and Lysine-tagged TMT (+304.207 Da). Percolator was used filter PSMs to 0.1%. Peptides were grouped using strict parsimony and only razor and unique peptides were used for protein level quantitation. Reporter ions were quantified from MS2 scans using an integration tolerance of 20 ppm with the most confident centroid setting. Only unique and razor (i.e., parsimonious) peptides were considered for quantification.

Data were deposited in the Pride database Project accession: PXD067627

### Weighted protein co-expression network analysis (WPCNA)

WPCNA analysis followed established WGCNA methodology and our previously published analytical pipeline (*32, 72, 73*). Proteomic Data Cleanup and Quality Control. Relative log_2_(abundance) data for either cortex or hippocampus from the TMT-MS quantitative summary by the search engine, normalized to within-batch global internal standard (GIS) sample abundance, was subjected to sample connectivity outlier removal. One outlier with sample connectivity more than 2.5 standard deviations below the mean was found for cortex and was removed. Proteins with 50 percent or more missing values were removed, resulting in matrices of 10,548 proteins x71 samples (for cortex) and 9,337 proteins x72 samples (for hippocampus). Nonparametric bootstrap regression was performed, modeling and removing covariance with TMT batch, and genetic background occurring within each protein in either the cortex or hippocampus protein relative log_2_(abundance) matrix. The removal of batch and background effects, and preservation of treatment group effects was confirmed using variancePartition R package (v 1.28.9) fitExtractVarPartModel() function.

Proteomics Systems Biology Pipeline. The data following quality control was input for coexpression analysis implemented in the R WGCNA package (v 1.72-1). Briefly, network scale free topology (SFT) was achieved by finding a suitable power to which the signed bicor correlation adjacency matrix could be raised to achieve a SFT R² of >0.80 (cortex: power 9, R²=0.832; hippocampus: power 9, R²=0.844) and achieving a median sample connectivity under 100. The WGCNA blockwiseModules() function was used to perform the multi-step network builds including adjacency matrix calculation, conversion to a topology overlap matrix (TOM), and protein pair dissimilarity metric calculations from 1-TOM, which serve as dendrogram distances allowing for dynamic hierarchical tree cutting to define clusters or modules of proteins. The module definitions on the dendrogram were further refined by partitioning about medoids, respecting the dendrogram. Other parameters for the function defining the initial network modules were a signed network type, mergeCutHeight of 0.07, a minimum module size of 20, and deepSplit=2, with maxBlockSize larger than the protein count, ensuring the full dendrogram was computed in a single block.

Cleanup of module membership enforcing that a protein’s module be reassigned if the bicor of a protein to a module eigenprotein is 0.10 bicor higher than the module originally assigned, followed by recalculation of module eigenproteins and iteration until resolution of all inconsistencies or 30 iterations, was performed as previously published (*73*). The final modules were then analyzed as a network of related modules, plotting (a) the protein relatedness dendrogram with module assignments and a bicor heatmap of each protein’s abundance profile to key traits; (b) the relatedness of module eigenproteins as a dendrogram; (c) bicor correlation of key traits to module eigenproteins; (d) a heatmap of eigenprotein values for each sample in the network, with relatedness dendrograms for both samples and eigenproteins (NMF R package aheatmap() function, v 0.26); (e) per each module, boxplots and scatterplots as appropriate for visualizing groupwise differences or continuous trait correlations, with appropriate statistics; (f) a heatmap of Fisher’s exact one-tailed enrichment for brain five cell type markers in each module generated using the open source function geneListFET() available from GitHub, leveraging the mouse marker gene product proteins we previously annotated as enriched via selective thresholding of abundances in purified bulk preparations from immunopanned populations performed in RNA-Seq (*74*) and proteomics (*75*) of the pure brain cell types; (g) ANOVA+Tukey post hoc test corrected volcano statistics calculated using an open source function parANOVA() and plotted using the plotVolc() function, both available from GitHub, which were also used to produce statistics and volcanoes for bicor correlation and significance of protein abundances to continuous variable traits; and (h) Gene Ontology (GO) enrichment of both modules and of ANOVA volcano comparison differentially abundant protein lists was performed and plotted using the GOparallel() open source function from GitHub, leveraging the July 2024 Bader Lab compilation of ontologies including GO (BP, MF, and CC), Broad Institute’s molecular signatures C2 database, Wikipathways, and Reactome.

Finally, the gene product proteins for 119 congruent RNA-protein markers, 33 CSF-brain markers, and 51 markers identified in the NULISA platform were checked for correlation to the module eigenproteins in both the cortex and hippocampus networks, and cross-region marker correlations were calculated using the WGCNA bicorAndPvalue() function and plotted in R. Linear modelling of cortex MeCP2 protein log2(abundance) as an outcome predicted by each of the 10,548 cortex proteins was performed to obtain effect size (β) and p value for each; the same analysis was performed for prediction of Bird score as outcome, and each set of p values was corrected to FDR using the Benjamini-Hochberg method. Volcano plots of β (x) vs. −log_10_(FDR) for each set of linear model runs (Mecp2 or Bird score) were plotted using the plotVolc(), and proteins with significant positive or negative association in each plot were subject to determination of ontology enrichment using GOparallel(). The 3 lists of interest described above were then intersected with the 10,548 proteins and only the common proteins kept for one additional volcano plot. The effect sizes of proteins from the three lists of interest were also plotted using ggplot() to compare Mecp2 association β (x) to Bird score effect size (y). Finally, proteins with a FDR-corrected significant effect size (either positive or negative) were counted as a fractions of each network module and plotted as a stacked bar plot across all modules in their module relatedness order. The bars were colored by the average β of those proteins within the module of the same sign (direction of association); the open-source function for this is DEXpercentStacked() in the parANOVA GitHub repository. All analyses were run in R v4.2.3.

https://github.com/edammer/GOparallel

https://github.com/edammer/CellTypeFET

https://github.com/edammer/parANOVA

https://download.baderlab.org/EM_Genesets/

### Feature selection by q value and Gene Product Annotations

DESeq2 generated coding and non-coding transcriptome normalized counts or proteome data were processed in Qlucore Omics Explorer 3.10.20 as log2 values. Data were fitted to a mean of 0 and variance of 1. We used multiple ANOVA or t-test with Benjamini-Hochberg multiple corrections to find q values under which genotypes were segregated using principal components 1 to 3 analysis and Euclidean distance of 0 between neighbors. Corrected p values obtained this way were within the range or exceeded the stringency of reported FDR corrected p values selected in the literature (*29, 76–79*).

Gene product annotations were analyzed using the v3.5.20250701 tool using the Multiple Gene List and Express analysis options and ENRICHR (*80–82*).

### Human Subjects

Clinical features of the two RTT cohorts and the neurotypical control group used in these studies have been previously described. The USA cohort was reported by Khwaja et al. (*37*) and the European cohort and the control group were described in publications by Naegelin et al and Zandl-Lang et al (*38, 83*). Briefly, all participants with RTT met established diagnostic criteria (3) and carried a *MECP2* pathogenic variant. Most of the females with RTT had a classic presentation, were in Hagberg’s stage III and approximately half of them had a severe clinical presentation. As shown in Fig. 7B, the USA cohort (median 6 years) was younger than the European (median 13 years) and the RTT group was overall younger (median 7 years) than the neurotypical control subjects (median 12 years). The referred studies were approved by the Institutional Review Board of Boston Children’s Hospital, the Ethics Committee of the Medical University of Graz, and the Northwestern and Central Switzerland ethics committee. Informed consent was obtained from the parent of each participant. CSF samples were received and remained deidentified for these studies.

### NULISA

Briefly, CSF samples were stored at −80°C until ready for use. Before the assay, samples were thawed and centrifuged at 10,000g for 10 min. 10 μL supernatant from the CSF sample was plated in 96-well plates and assayed with the NULISAseq CNS Panel 120, targeting mostly neurodegenerative disease-related protein markers and inflammation and immune response-related cytokines and chemokines. Mouse plasma cytokines were assayed with the NULISAseq CNS Panel 220 using 10 μL of biofluid.

All steps from immunocomplex formation with the paired set of oligo-conjugated antibodies, first capture with oligo-dT beads, release, second capture with streptavidin beads and ligation were performed on the automated Alamar ARGO^TM^ _prototype system. The library of DNA reporters containing unique target-specific molecular identifiers (TMI) and sample-specific molecular identifiers (SMI) was pooled, amplified by PCR, purified and sequenced on the Illumina NextSeq 2000 system. Data were processed using discriminant partial least squares regression analysis (*43, 44*) to identify analytes discriminating genotypes and in a parallel dataset was thresholded by p value.

### Differential Expression and Partial Least Squares Discriminant Analysis

Differentially expressed proteins (DEPs) were identified between Rett syndrome and neurotypical individuals using the *lmNULISAseq* function in the NULISAseqR package (v1.2.0). This function utilizes a linear regression model to identify DEPs. Significant DEPs were those with FDR adjusted p-value less than 0.05. Partial least squares discriminant analysis (PLS-DA) was performed using the *opls* function in the ropls package (v1.36.0) in R, after the data were z-scored. Variable importance in projection (VIP) for each protein was determined using the *getVipVn* function in ropls. A VIP value greater than 1 was considered to be important. Standard deviation of LV1 loading for each protein was determined using a leave-one our cross validation (LOOCV) approach, whereby the PLS-DA projection is recalculated with one sample removed and repeated iteratively across all samples. The output of PLS-DA was rotated to best capture group-wise differences.

### Immunoprecipitation of VGF from *MECP2* KO/Control SHSY5Y conditioned media

SHSY5Y Control and *MECP2* KO cells were developed by Synthego and described by Zlatic et al 2023. Cells were cultured in DMEM (Corning 10-013-CV) with 10% FBS (VWR 97068-085) at 37°C, 5% CO^2^. Equal volumes of media, conditioned by equal cell counts for 12 hours, was collected and clarified of cellular debris at 21,000 RCF for 20 minutes at 4C. Immunoprecipitation was performed as described previously (*84*). Briefly, Dynabeads (Thermo 11203D) loaded with 0.2ug of anti-VGF (Thermo PA5-63081) were incubated overnight at 4C with clarified media containing Complete antiproteases (Roche 11245200), 0.5% Triton-X 100, and 1x buffer A (150 mM NaCl, 10 mM HEPES, 1 mM ethylene glycol-bis(β-aminoethylether)-N,N,N′,N′-tetraacetic acid (EGTA), and 0.1 mM MgCl2, pH 7.4).

### Mouse Cortex Lysis

Cortex collected from Control and *Mecp2* KO mice (N=6 per genotype) was lysed in 0.5% Triton-X 100, and 1x buffer A containing Complete antiprotease by sonication and kept on ice for 30 minutes. Lysed tissue was clarified at 16,100 × *g* for 15 min, and the supernatant was recovered. Protein concentration of the clarified lysate was determined by Bradford Assay (Bio-Rad, 5000006.)

### SDS-PAGE and Western Blot

Washed IP beads, conditioned media inputs, and mouse cortex lysis were treated with denaturing Laemmli buffer, heated for 5min at 75C, run on 4-20% Criterion gels (Bio-Rad, 5671094) and Western blotted onto PVDF membrane using the semi-dry method. Membranes were probed with anti-VGF (Thermo PA5-63081), AACS (Proteintech, 13815-1-AP) Calb1 (Proteintech, 14479-1-AP), AGXT2L1/Etnppl (Proteintech, 83984-6-RR), PTN (Cell Signaling, 54934T), Got1 (Proteintech, 14886-1-AP), Actb (Sigma, A5441), Mecp2 (Invitrogen PA1-888) and anti-rabbit or mouse HRP secondary antibodies (Thermo Fisher Scientific, A10668 and G-21234) and detected with Western Lightning Plus ECL reagent (PerkinElmer NEL105001EA) on GE Healthcare Hyperfilm ECL (28906839). Conditioned media SDS-PAGE was also stained with Coomassie Blue (BioRad 1610803).

### Targeted Metabolomics

Flash-frozen brain tissues combined from both right and left hemispheres were ground using a liquid nitrogen-chilled porcelain mortar and pestle. 1 ml of ice-cold 4:4:2 (methanol:acetonitrile:water) containing 0.1% formic acid and 25 µM of 13C3 alanine internal standard was added to 25 mg of ground tissue powder on ice and briefly vortexed to mix. At 3 minutes of incubation, 15% ammonium bicarbonate was added to neutralize the pH. Samples were then sonicated on ice for 3 pulses (5 sec on/5 sec off) at a power of 15 (Microson ultrasonics cell disruptor XL). To ensure complete tissue lysis, samples underwent 3 freeze/thaw cycles in liquid nitrogen and ice. The samples were then left on ice to maximize protein precipitation, followed by centrifugation at 21,000 x g at 4 °C for 30 min, and the supernatant was concentrated to dryness using a Savant SpeedVac Plus vacuum concentrator. Dried samples were resuspended in 120 µl of ice-cold 60:40 acetonitrile:water, briefly vortexed, and sonicated in an ice bath for 5 min to solubilize metabolites. Finally, samples were centrifuged at 21,000 x g at 4 °C for 30 min, and 100 µl of supernatant was transferred to an LCMS vial for mass spectrometric analysis. 5 μl of this sample was injected onto a HILIC-Z column (Agilent Technologies), and metabolites were measured using an Agilent 6546 QTOF mass spectrometer coupled with an Agilent 1290 Infinity II UHPLC system (Agilent Technologies). The column temperature was maintained at 15 °C, and the autosampler was kept at 4 °C. Mobile phase A consisted of 20 mM Ammonium Acetate, pH = 9.3 with 5 μM Medronic acid, and mobile phase B was 100% acetonitrile. The gradient, run at a flow rate of 0.4 ml/min, was as follows: 0 min, 90% B; 1 min, 90% B; 8 min, 78% B; 12 min, 60% B; 15 min, 10% B; 18 min, 10% B; and 19-23 min, 90% B. The mass spectrometry data were collected in negative and positive modes over an m/z range of 20-1100 at 1 spectrum/sec, with the following instrument parameters: gas temperature 225 °C, drying gas flow 9 l/min, nebulizer pressure 10 psi, sheath gas temperature 375 °C, sheath gas flow 12 l/min, VCap 3000 V (negative mode) or 3500 V (positive mode), nozzle voltage 500 V, fragmentor 100 V, and skimmer 45 V. A pooled quality control sample consisting of 5 µl from each experimental sample was injected after every 5 experimental samples and used to detect metabolites that degraded in the instrument during run and were not included in the analysis. Data were analyzed using MassHunter Qualitative Analysis 10 and MassHunter Quantitative Analysis 11 (Agilent Technologies).

### NAD+/NADH Assay

The Promega NAD/NADH Glo^TM^ Assay with modifications based on Wang et al 2024 was used to measure NAD/NADH levels from Control and *Mecp2* KO mouse cortex (*85*). Briefly, tissue was weighed and sonicated in PBS then clarified by centrifugation at 13,000RPM for 10 minutes. Clarified lysis was passed through a 10KDa spin column (Millipore MRCPRT010). Eluent was processed for isolation and detection of NAD and NADH as per Manufacturers protocol. Luminescence was detected with a BioTek Synergy HT microplate reader running Gen6 software. Luminescence was reduced by the average blank value and normalized to Control mouse tissue values.

### Statistical Analyses

Unless otherwise indicated, all statistics were performed with either Prism Version 10.5.0 (673), Estimation Stats (estimationstats.com/#/), or Qlucore Omics Explorer 3.10.20. Data were tested for normality to apply the proper test. N values reported in each figure correspond to biological replicates, individual animals, or subjects. All p vales reported are two-tailed. Except for figure 6, no outlier identification was performed.

## List Supplementary Materials

**Supplementary Table I. Data supporting Figures 1-4 and 6-8 and their supplements**

10.6084/m9.figshare.33005975

**Supplementary Table II. Data supporting Weighted protein co-expression network analysis (WPCNA) and their supplements**

10.6084/m9.figshare.33005975

## Acknowledgments

The authors would like to express their gratitude to all the members of the Rett community for their support and advocacy and members of the Faundez Laboratory for their comments and suggestions. Maria Olga Gonzalez provided NAD and NADH standards for VF.

## Author contributions

Conceptualization: VF, RC, SC

Methodology: SZ, AC, ED, DD, JS, KKEG, AG, BT, LW, AP, VF

Investigation: SZ, AC, ED, DD, JS, KKEG, AG, BT, LW, AP, VF

Human Samples: MZ-L, BP, LA, WK

Visualization: VF, ED

Funding acquisition: VF, SC

Project administration: VF

Supervision: VF, SC

Writing – original draft: VF

Writing – review & editing: SZ, AC, ED, DD, JS, KKEG, AG, BT, LW, AP, BP, RC, SC, VF

## Data and materials availability

All data are available in the main text or the supplementary materials. Accession information for Pride and GEO files can be found in materials and methods.

## Supplemental Materials for

**Figure 1S1.**
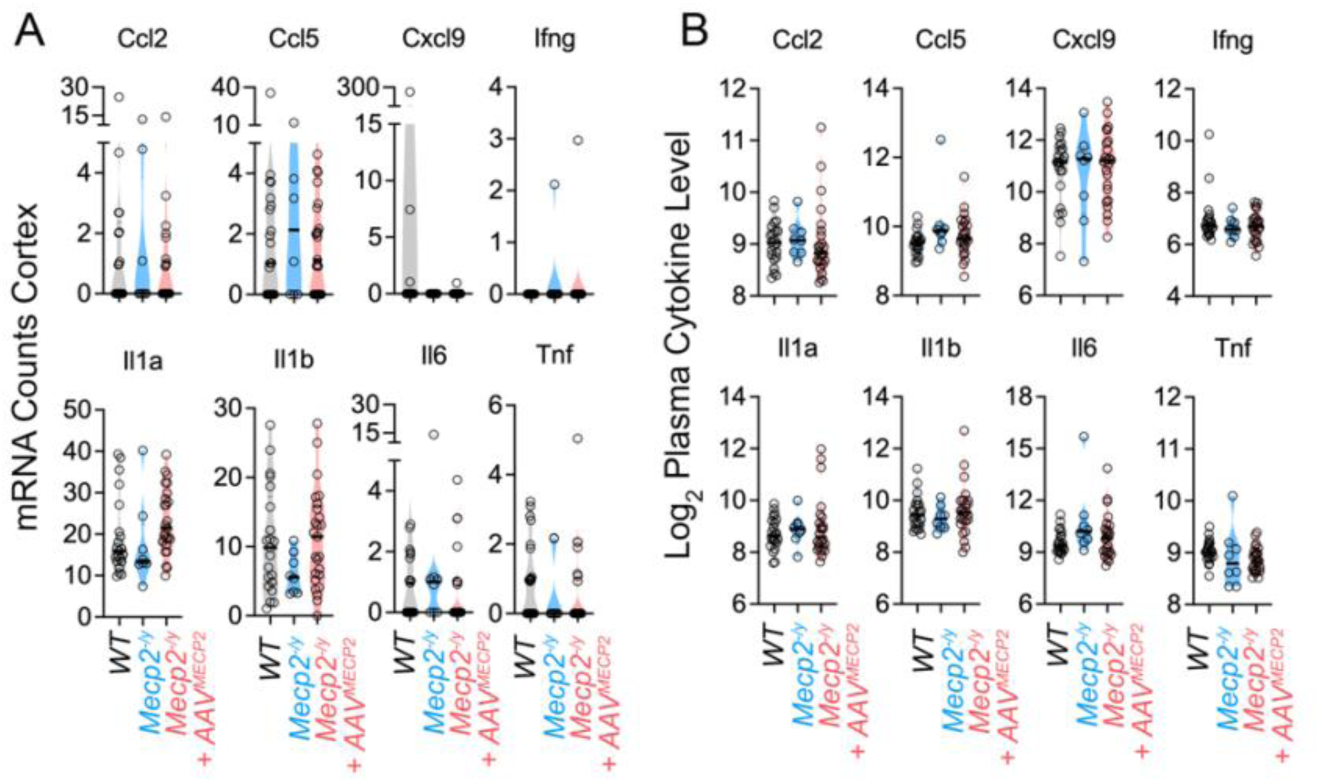
Cytokine Brain mRNA and Plasma Levels in Wild type, *Mecp2^-/y^* and the *Mecp2* gene therapy-treated Animals. Cortical cytokines mRNA levels measured by RNAseq (A) and plasma levels (B) determined by NULISA in the mouse cohort described in Fig. 1 and analyzed by multiomics in Figures 1 to 5. One way ANOVA reveals no statistical differences across experimental groups in both assays.

**Figure 2S1.**
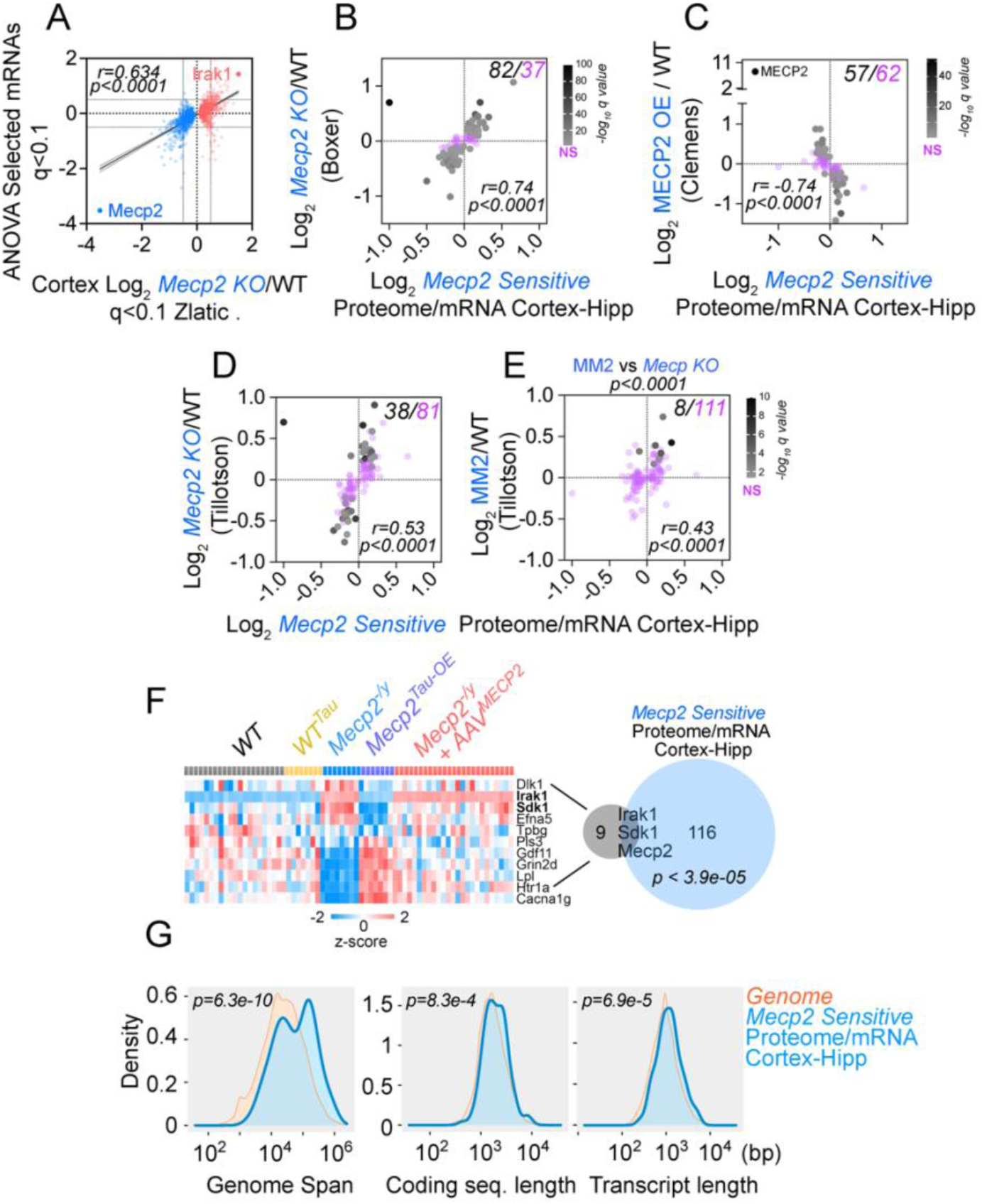
Correlations between *Mecp2^-/y^* Cortex Transcriptomes and the *Mecp2* mutant and gene therapy-sensitive mRNA-Protein Pairs. A. Multiple ANOVA q<0.1-defined mRNAs from *Mecp2* mutants and gene therapy animals (see Fig. 2C) were correlated to the cortical transcriptome by Zlatic et al (2023) thresholded at the same q value. B-E. The 119 Mecp2 mutant- and gene therapy-sensitive mRNA-Protein Pairs (See Figure 3G) correlate with the mRNA identified in *Mecp2^-/y^* cortices in two studies -Boxer et al, Tillotson et al. and a mouse model overexpressing MECP2 by Clemens et al (C) Pearson correlation coefficient. D and E depict RNAseq results comparing a *Mecp2* null mouse and a *Mecp2* mutant mouse carrying a DNA binding domain that does not bind to mCAC sequences (MM2) by Tillotson et al. MM2 vs *Mecp2* KO compares the number of significantly changed transcripts in both animal models (*p*<0.0001, two-sided exact binomial McNemar test). A-E number on the upper right corner show significant (black) and non-significant mRNA (purple) in the published datasets overlapping with the 119 *Mecp2* mutant- and gene therapy-sensitive mRNA-Protein Pairs. F. Twelve core *Mecp2*-sensitive transcripts defined by Li et al. (*11*) display coherent profiles in the cortex multiple ANOVA q<0.1-defined mRNAs. Three transcripts are shared with the 119 *Mecp2* mutant- and gene therapy-sensitive mRNA-Protein Pairs (See Figure 3G). D. The 119 *Mecp2* mutant- and gene therapy-sensitive mRNA-Protein pairs are enriched in long genes and transcripts. T-test compares the difference between the 119 proteins and the other protein-coding genes on the genome (ShinyGo 0.82(*64*))

**Figure 2S2.**
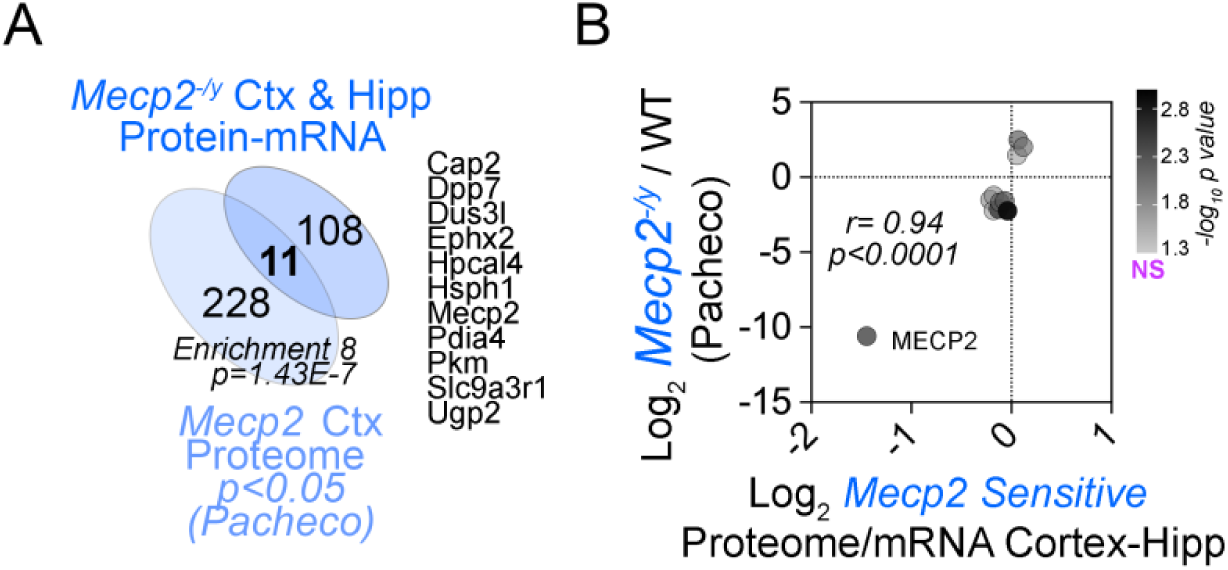
Correlation between 119 *Mep2* sensitive mRNA-Protein Pairs with a Published Proteome Dataset. A. Venn diagram depicts the overlap between the 119 *Mecp2* mutant and gene therapy-sensitive mRNA-Protein Pairs (see Fig. 3G), and the and the significant proteome dataset by Pacheco et al. Overlap represents an enrichment of 8 fold (p < 1.43e-7, Exact hypergeometric probability). B. The 119 *Mecp2* mutant- and gene therapy-sensitive mRNA-Protein Pairs (See Figure 3G) correlate with the 11 proteins shared with Pacheco et al. Pearson correlation coefficient.

**Figure 3S1.**
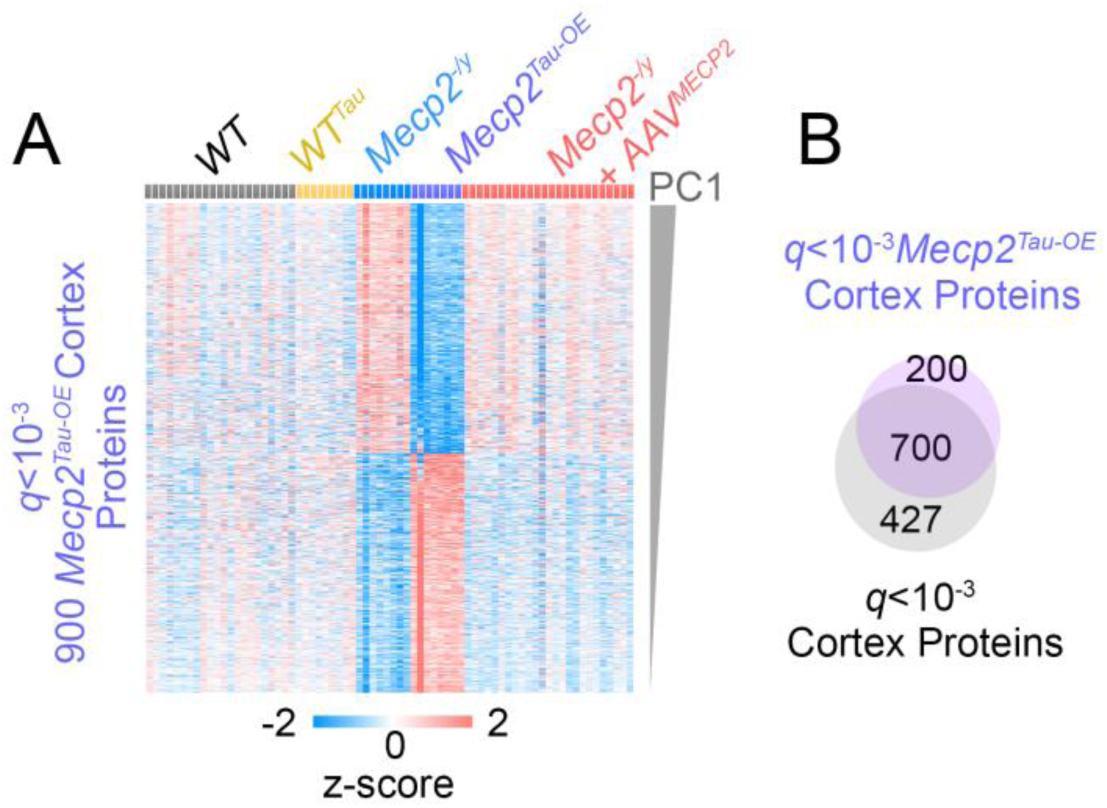
q-defined *Mecp2^Tau-OE^*Cortex Proteome Identifies Proteins Altered by Mecp2 Overexpression. A. *Mecp2^Tau-OE^* cortical Z-scored heat map of proteomes selected by FDR-corrected t-test comparing *Mecp2^Tau-OE^* to all other groups combined q<0.001. Rows are organized by analyte contribution to PC1. B. Venn diagram depicts the overlap between the multiple ANOVA q-defined cortex proteome and the t-test q-defined *Mecp2^Tau-OE^* cortex proteome. There are 200 proteins whose levels are sensitive only to the *Mecp2^Tau-OE^* allele. Processed data used to build figures can be found in supplementary table I.

**Figure 3S2.**
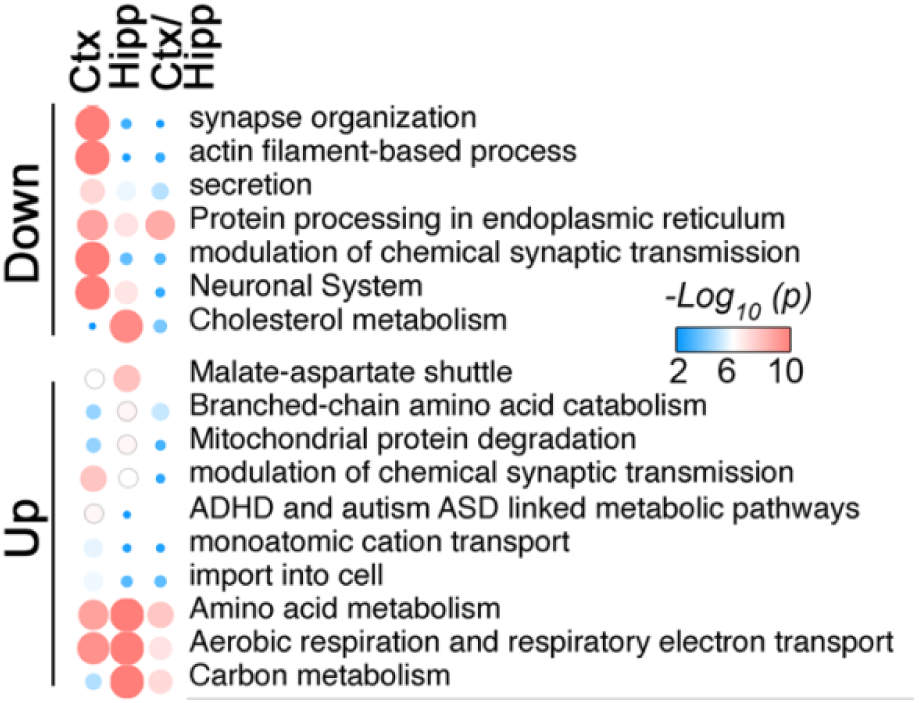
Shared gene ontologies among up and down-regulated proteins comparing cortex (584 proteins), hippocampus (198 proteins), and the 119 mRNA-Protein Pairs. Metascape shared gene ontologies among up and down-regulated proteins comparing cortex (from the 584 proteins selected in Fig. 3E), hippocampus (from the 198 proteins selected in Fig. 3F), and the 119 proteins in Fig. 3G. Symbol size and color are proportional to p value. Processed data used to build figures can be found in supplementary table I.

**Figure 5S1.**
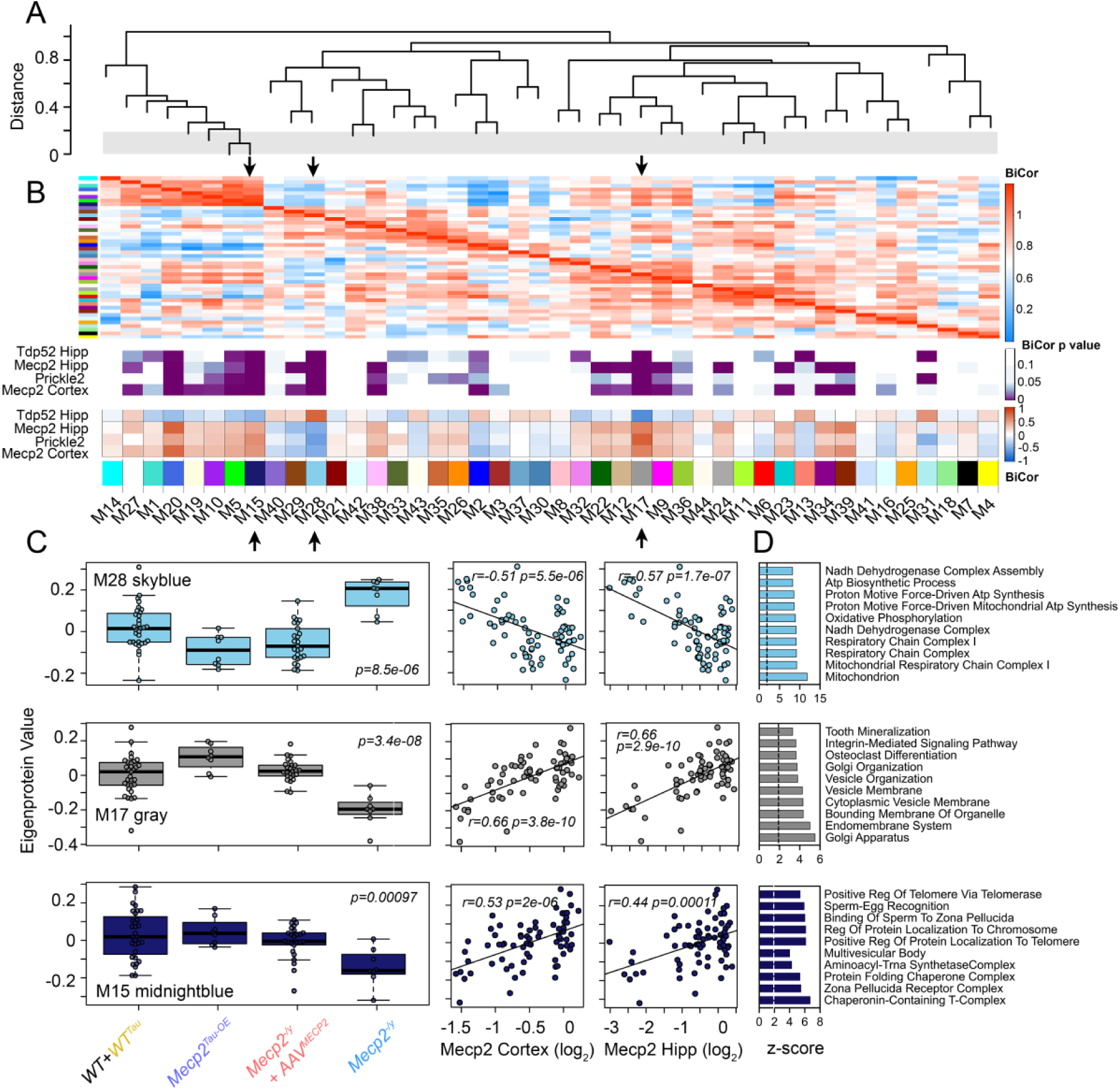
Hippocampal Protein Coexpression Network of *Mecp2* Mouse Models without and with MECP2 Gene Therapy. A. Weighted protein co-expression network analysis (WPCNA) grouped proteins into distinct protein modules (M1–M44) clustered to assess module relatedness based on correlation of protein co-expression eigenproteins. B. Biweight midcorrelation (BiCor) analysis of module eigenproteins with either Mecp2, Prickle2 and Tdp52 protein levels, and the aggregated behavioral score as traits. Purple heat map shows significance of association of each trait to a module using Benjamini-Hochberg FDR-corrected Student correlation p value. Red-blue heat map indicates strength of association where red represents positive correlation and blue indicates negative correlation of eigenproteins to each module. Module-trait correlations were performed using biweight midcorrelation (bicor). C. Changes in eigenprotein abundance in salient modules indicated by arrows across *Mecp2* mouse models without and with *MECP2* Gene Therapy. multiple ANOVA with Tukey post hoc correction was used to assess the statistical significance of abundance changes. Eigenprotein correlation with indicated modules for each subject using Mecp2 content in two brain regions and the aggregated score as traits (Biweight midcorrelation). Selected modules with significant trait correlations were analyzed by gene ontology (GO) analysis, dotted line marks Z-score of 2. Processed data used to build figures can be found in supplementary table II.

**Figure 5S2.**
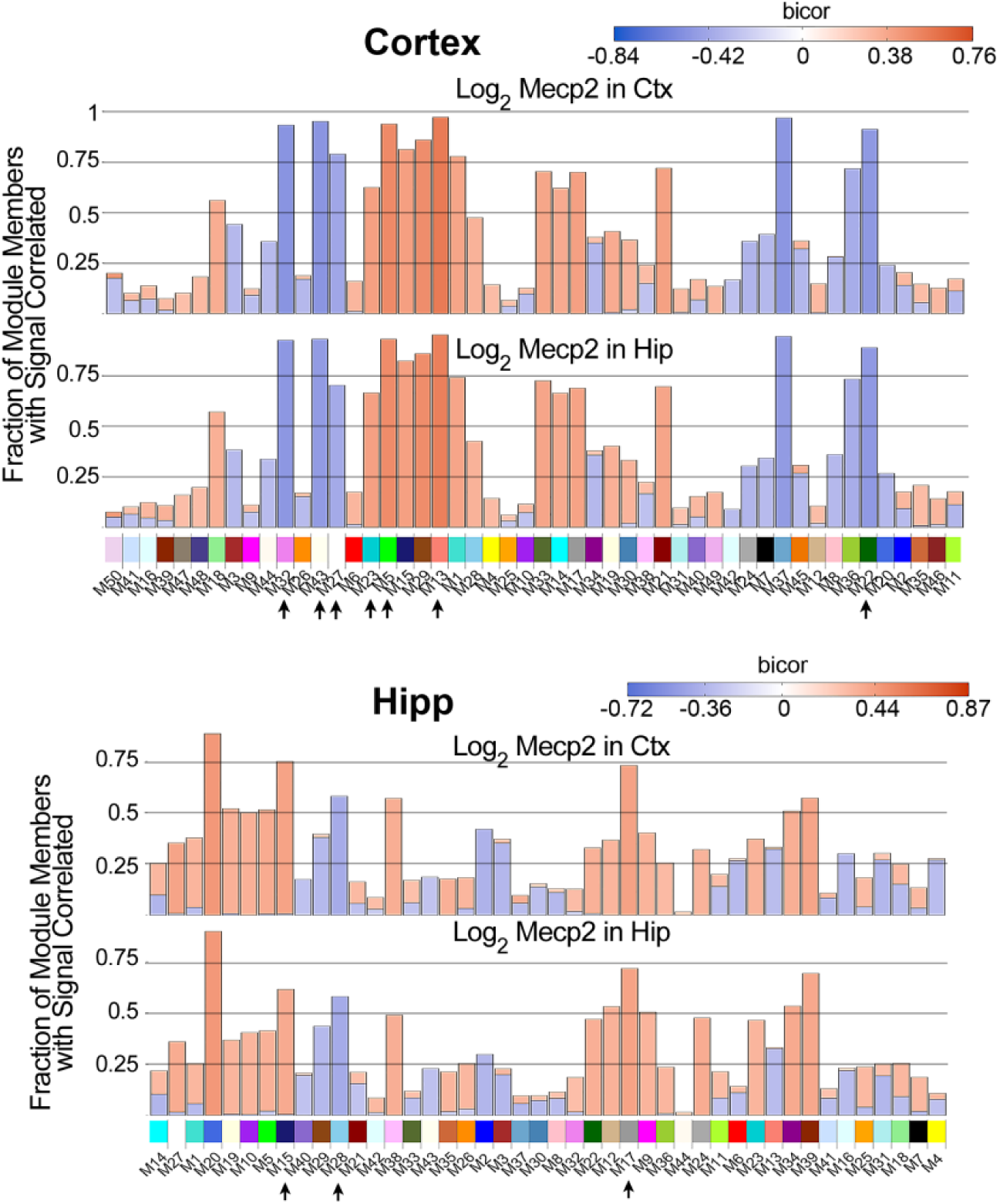
Fraction of Differentially Expressed Proteins in the Coexpression Network of *Mecp2* Mouse Models without and with *MECP2* Gene Therapy. Cortex and hippocampal networks of differentially expressed proteins were assessed for the fraction of module member proteins correlating with the module eigenprotein value. Blue and red correspond to negative and positive Biweight midcorrelation r values. Processed data used to build figures can be found in supplementary table II.

**Figure 7S1.**
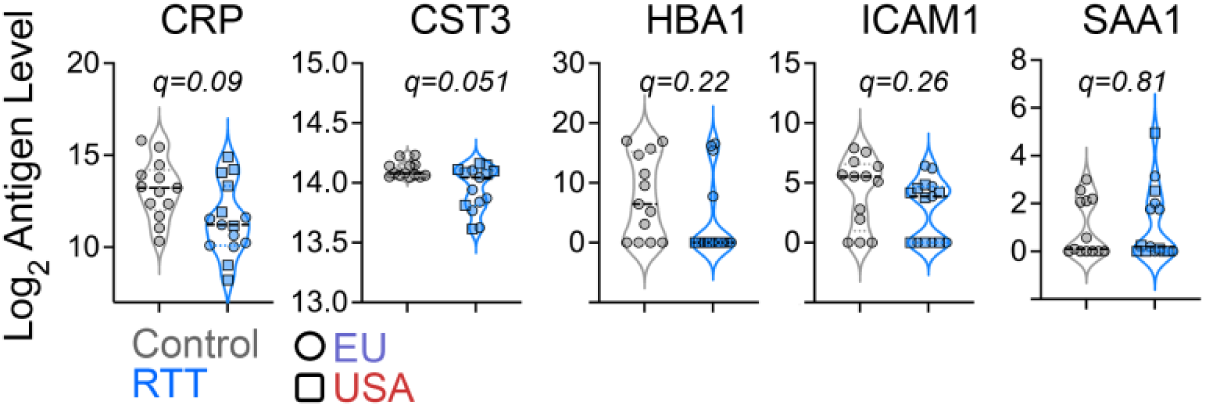
Composition of Neurotypical and Rett Syndrome Subjects Cerebrospinal Fluid Measured by NUcleic acid Linked Immuno-Sandwich Assay (NULISA). NULISA quantification of abundant blood proteins in neurotypical and RTT samples. T-test followed by Benjamini-Hochberg FDR correction.

**Figure 7S2.**
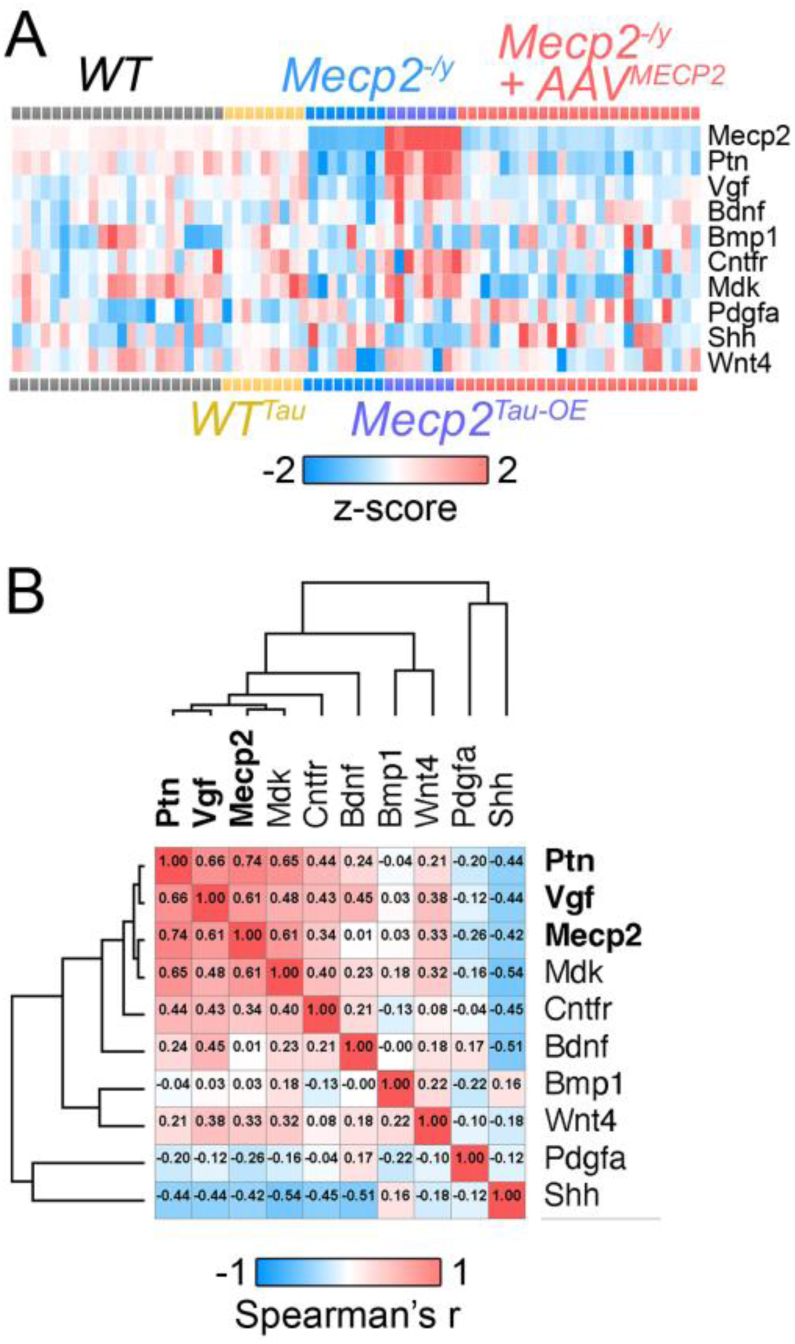
Differential Expression of Growth Factors in the Cortical Proteomes of *Mecp2* Mutant Mice and their Restoration by Gene Therapy. A.Z-scored heat map of growth factors quantified in the proteomes of *Mecp2* gain- and loss-of function mutation mouse cortex and the effect by AAV9-RTT271 therapy. B. Similarity matrix of analytes in A. Numbers depict analyte correlations. Processed data used to build figures can be found in supplementary table I.

**Figure 7S3.**
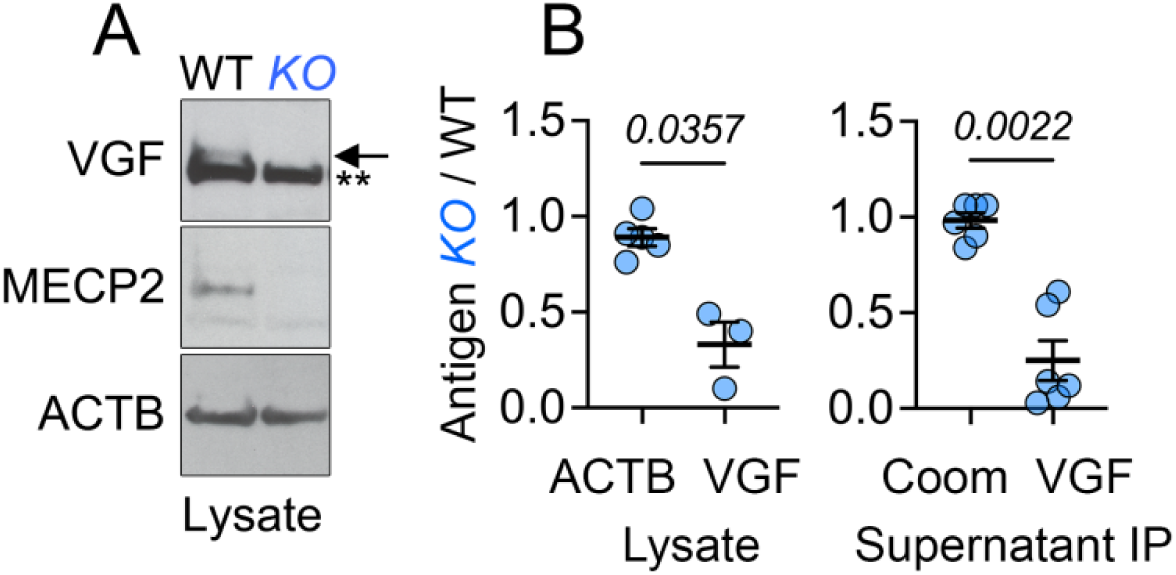
VGF Expression is Reduced in *MECP2*-null Human Neuroblastoma Cells. A. Wild-type and *MECP2*-KO SH-SY5Y neuroblastoma cell lysate probed with antibodies against VGF (arrow), MeCP2, and beta actin (ACTB). Double asterisks mark non-specific band. Conditioned media show decreased levels of VGF is Figure 6I. B. Quantifications of blots in Fig. 6H and in panel A. Mean ± SEM. P value calculated with two-tailed t-test.

**Figure 8S1.**
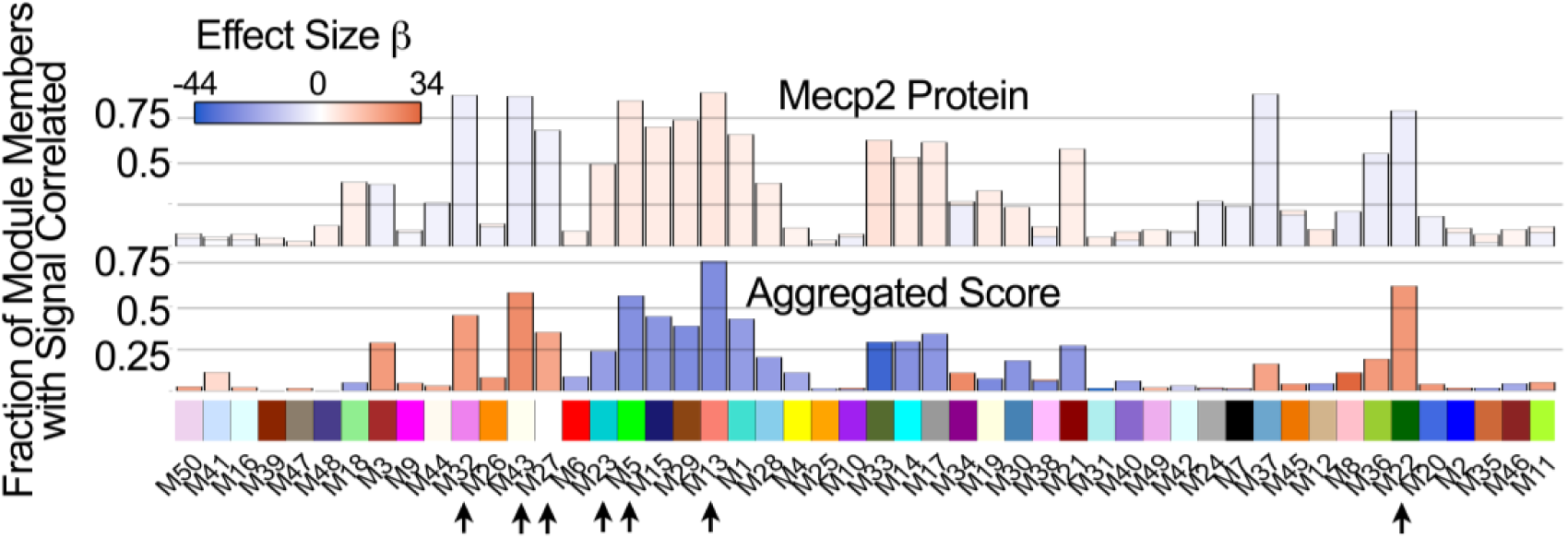
Fraction of Differentially Expressed Proteins in the Brain Mecp2 Coexpression Network after Analyte Selection by their Effect Size on MeCP2 and Aggregated Score Traits. Cortex and hippocampal networks of differentially expressed proteins were assessed for the fraction of module member proteins correlating with the module eigenprotein value. The proteins used in the analysis correspond to the sum of the three datasets used to calculate effect size in Fig. 7A. Blue and red correspond to negative and positive beta scores for Mecp2 protein content and aggregated clinical score as traits. Processed data used to build figures can be found in supplementary table II.

**Figure 8S2.**
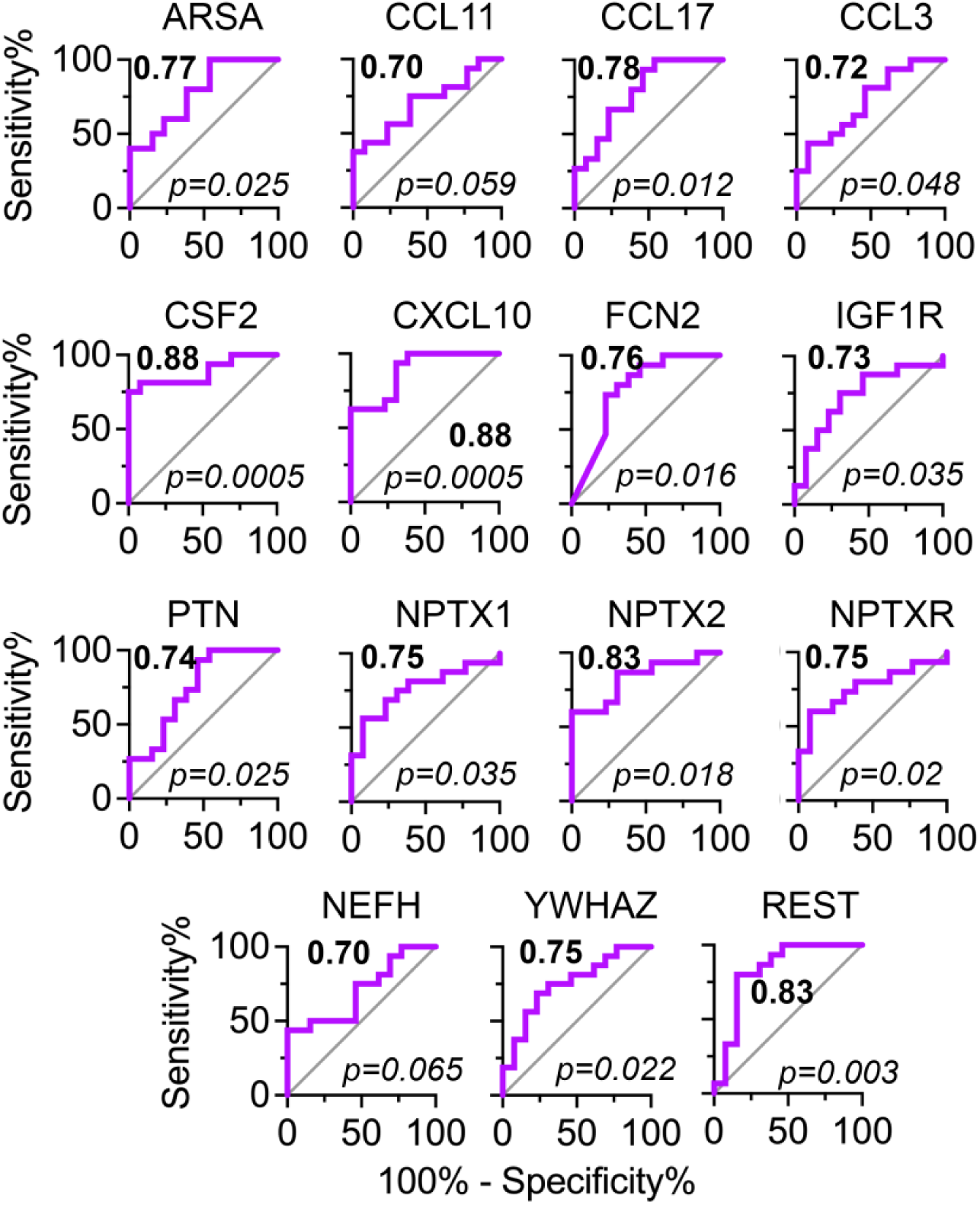
Discriminant Power of Proteins Curated by *Mecp2* Genotypes, *MECP2* Gene Therapy, and Rett Syndrome Diagnosis. Discrimination between neurotypic and RTT CSF by selected analytes determined by the area under the curve (AUC, bold number) in Receiver Operating Characteristic (ROC) analysis.

